# Water stress-associated isolation barriers between two sympatric oak species

**DOI:** 10.1101/2021.09.16.460585

**Authors:** Grégoire Le Provost, Benjamin Brachi, Isabelle Lesur, Céline Lalanne, Karine Labadie, Jean-Marc Aury, Corinne Da Silva, Dragos Postolache, Thibault Leroy, Christophe Plomion

## Abstract

Drought and waterlogging impede tree growth and may even lead to tree death. With climate change, these environmental factors are a growing source of concern, particularly for temperate forests. Oaks, an emblematic group of tree species, have evolved a range of adaptations to cope with these constraints. The two most widely distributed European species — pedunculate oak (PO) and sessile oaks (SO) — have overlapping ranges, but are highly constrained locally by soil water content variation. These differences in local ecological requirements provide a powerful biological model for studying the role of ecological barriers in speciation. We used an experimental set-up mimicking the ecological preferences of these species, in which seedlings were subjected to waterlogging and drought. We studied gene expression in roots by RNA-seq and identified genes differentially expressed between treatments with different outcomes depending on species. These “species x environment”-responsive genes revealed adaptive molecular strategies involving adventitious and lateral root formation, aerenchyma formation in PO, and osmoregulation and ABA regulation in SO. With this experimental design, we also identified genes with expression profiles presenting a “species” effect regardless of imposed constraints with important roles in intrinsic reproductive barriers. Finally, we compared our findings with those for a genome scan of species divergence and found that the candidate genes were enriched in highly differentiated SNPs. This suggests that many of the genes involved in the contrasting transcriptomic responses are subject to natural selection and that gene regulation helps to maintain these two different oak species in sympatry.

## Introduction

One of the major goals in evolutionary biology is understanding the genetic architecture of speciation, the number of genes involved in the evolution of reproductive barriers, their effect sizes and functions (Campbell et al, 2018)). There is now a clear consensus among evolutionary biologists about the continuity of the speciation process, including the gradual accumulation of barriers to gene flow in geographically isolated populations (Coyne & Orr, 2004). However, the long-standing debate about the contribution of local adaptation to the initiation and development of reproductive isolation between non-geographically isolated populations remains unresolved. This question is particularly difficult to answer given that most species are known to have experienced successive periods of isolation and contact in the past (Bierne et al., 2011). It remains possible to identify loci that have contributed to reproductive isolation (referred to hereafter as barriers to gene flow) in pairs of closely related species that are still exchanging genes, and to formulate functional hypotheses about the ecological or non-ecological (intrinsic) nature of reproductive barriers, by considering gene annotations. It is, indeed, possible to identify ecological and non-ecological barriers to gene flow by scanning genomes for regions with high levels of relative divergence. Many scientific articles published over the last decade have identified candidate reproductive barrier genes through analyses of whole-genome sequence data from closely related species (reviewed by Ravinet et al., 2017; Wolf et al., 2010).

As a complementary approach to classical studies aiming to analyze the level of sequence divergence between sister species (reviewed by Storz, 2005), studies of the relationships between divergences in gene expression and species persistence have also yielded interesting results. Fay and Wittkopp (2008) suggested that variations in transcription levels have a tremendous potential for generating evolutionary novelty that can then be harnessed by natural selection. This approach is based on the detection of genes displaying significant differential expression (significantly differentially expressed genes or DEGs) between two or more focal species and searching for (i) genes displaying “species-specific” differential expression, regardless of the environmental conditions, or (ii) “species-specific” DEGs expressed in particular environments. The first group would be expected to contain intrinsic reproductive isolation barriers between sister species, whereas the second, containing genes displaying significant species-by-environment interactions, would be expected to include important genes underlying the strategies of the species concerned for coping with the challenging environmental conditions encountered in its ecological niche. The contribution of these ecological barriers to overall reproductive isolation (sometimes referred to as ecological speciation) remains a matter of debate. However, these specific contributions can be deciphered by identifying genes in an experimental set-up mimicking the different environmental requirements contributing to the ecological preferences of adapted species and, consequently, to overall reproductive isolation.

Most of the transcriptomic research studies on speciation performed to date involved the use of animal models, including *Littorina saxatilis* marine snails (Martínez-Fernández et al., 2010), the lake whitefish (*Coregonus clupeaformis*; Renaut et al. 2010, Jeukens et al., 2010) and Darwin’s finches (Abzhanov et al., 2006). In plants, the role of diverging gene expression in speciation has been studied mostly in hybrid systems. For example, in *Senecio* species, some studies have reported profound changes in gene expression in new hybrids, favoring the survival of these plants in new environments not accessible to their parents (Chapman et al.; 2013, Hegarty et al., 2008). In sunflower, Lai et al., (2006) found that environmental pre-adaptation was driven mostly by expression divergence (reviewed by Mack & Nachman, 2017). However, in forest trees, very few such studies have been performed (reviewed by (Lexer et al., (2004)).

European white oaks are an excellent model for investigating such divergence in expression. These sister species diverged recently (<10 million years ago (mya), Hubert et al., 2014; Manos & Hipp, 2021) and have probably experienced a series of periods of allopatric isolation and contact associated with glacial retreats and postglacial recolonizations (Leroy et al., 2017; 2020). Within this species complex, the speciation of sessile oak (*Quercus petraea* Matt. Liebl., hereafter SO) and pedunculate oak (*Quercus robur* L. hereafter PO) has been studied in detail (reviewed in (Kremer & Hipp, 2020). In “On the origin of species by means of natural selection, or preservation of favoured races in the struggle for life”, Darwin himself — in an attempt to define species and speciation introduced pedunculate and sessile oaks as an example illustrating the difficulties of clearly defining different species on the basis of very closely related but different life forms. Indeed, PO and SO are known to hybridize (Lepais & Gerber, 2011) and their distinction on the basis of morphological criteria is not straightforward, to the extent that some evolutionary biologists have described them as “botanical horrors” (Rieseberg, Wood, & Baack, 2006). That said, they do have known different ecological preferences imposed largely by environmental factors, such as light, soil composition and humidity (Epron & Dreyer, 1990), although there is some overlap between their habitats (Leroy et al., 2020). In brief, SO is more tolerant of water shortage (Bréda, et al., 1993) and has a higher water-use efficiency than PO (Ponton et al, 2002). It is therefore better able to survive both chronic and extreme droughts (Bréda & Badeau, 2008). The severe heat waves and prolonged rainfall deficit observed in mainland France in 1976 and 2003 provide a clear illustration of the long-term consequences of such events in terms of oak tree decline in France and, more broadly, in Europe (Breda et al., 2006; van der Werf etal., 2007). Remarkably, these events had different effects on PO and SO (Thomas & Hartmann, 1996), with SO displaying higher growth rates during these dry years (Becker et al., 1994; Bréda & Badeau, 2008; Friedrichs et al., 2009; Lebourgeois, 2006) and PO populations having a higher mortality rate (Durand et al., 1983; Lebourgeois et al., 2015). Similarly, Truffaut et al. (2017) provided evidence for changes in species occupancy at a fine spatiotemporal scale in a mixed stand of these two species, with an expansion of the SO population attributed to differences in survival after the repeated drought events observed in the study area (west of France) over the last three decades. Conversely, PO can tolerate long periods of waterlogging and is generally found in moist habitats. Based on these observations and their consequences in terms of the adaptation of oak forest management to a warming climate, research has focused on characterizing the differences between SO and PO, with the aim of incorporating the knowledge obtained into management decisions. Differences between SO and PO have mostly been addressed at the phenotypic level (Parelle et al. (2007), for traits relating to hypoxia tolerance (Dupouey & Badeau, 1993; Kremer et al., 2002), leaf and fruit morphology and water use efficiency (Ponton et al., 2002). Leroy *et al*. (2020) recently addressed this question from a population genomics perspective, making use of the the oak genome sequence (Plomion et al. 2018) to perform genome scans for divergence between the four main European white oak species, including PO and SO. They identified candidate genes for reproductive isolation between species, including a large number of transcription factor genes. This study was particularly informative about the reproductive barriers between temperate (*Q. robur* & *Q. petraea*) and Mediterranean species (*Q. pyrenaica* & *Q. pubescens*), but shed little light on the genes contributing to the specific reproductive barriers between *Q. robur* and *Q. petraea*. Only a few studies have investigated the differences in gene expression patterns between SO and PO. Porth et al. (2005) provided a first insight into species differences in mRNA levels between these two species in conditions of water deficit. They detected an upregulation of osmotic adjustment-related genes in SO. In a study based on suppressive subtractive hybridization, Le Provost et al. (2012) provided further insight into the genes potentially involved in the adaptation of PO to waterlogging. In particular, they identified genes with expression levels displaying strong species-by-environment interactions. These observations suggested that PO had evolved specific strategies to cope with waterlogging. They also showed that tolerance to waterlogging was driven by a rapid switch of PO metabolism to the fermentative pathway in hypoxic conditions, a strategy that could alleviate energy loss caused by oxygen deprivation.

Our objective here was to investigated in more detail the contribution of gene expression to the maintenance of these two species in sympatry. We subjected PO and SO seedlings to environmental conditions mimicking the ecological constraints (i.e. an excess or deficit of water) to which they are adapted in natural conditions. We then harvested root tips, quantified gene expression by RNA-seq and identified the main factor (species, environment, their interaction) accounting for differential gene expression. We hypothesized that genes displaying a significant “species x environment” interaction effect would shed light on the different molecular strategies that had evolved in each species for adaptation to the most frequently encountered type of water stress, whereas genes displaying a significant “species” effect regardless of environmental conditions (i.e. always upregulated in either PO or SO) would highlight important functions underlying intrinsic barriers between these two species. We also combined our expression data with data from a whole-genome scan. We showed that the two categories of candidate genes described above were enriched in SNPs displaying a high level of genetic differentiation between the two species, confirming that these candidate genes play an important role in building extrinsic and intrinsic reproductive barriers between these two co-occurring species.

Overall, this study suggests that genes involved in molecular plasticity to the environmental conditions faced by these two species in their respective ecological niches are targets of divergent selection between the two species. This finding improves our understanding of the genetic basis of ecological speciation in European white oaks, and the identification of these candidate barrier loci is of prime importance for the design of sustainable oak forest management strategies to guide the trajectory of these species more effectively in the current context of climate change.

## Materials and Methods

### Plant material

The plant material used here was described in a previous study by Le Provost *et al*. (2016). Briefly, we sampled half-sib progenies from three unrelated mother trees for both SO (located in “Foret Domaniale” of Laveyron, latitude 43°45′49″N, longitude 0°13′11″W) and PO (from “Ychoux forest”, latitude 44°33′33″N, longitude −0°96′66″W). Before germination, we confirmed the species status of the mother trees with the diagnostic SNP markers described by Guichoux *et al*. (2011). We harvested the acorns in the fall of 2013 and sowed them in a 1:1 mixture of peat and sand in 0.2 L pots. After germination, we transferred the seedlings to a greenhouse (16-hour photoperiod, 25°C during the day and 20°C at night) for five weeks, until they had three fully developed leaves.

### Experimental design and measurement of physiological traits

Sixty homogeneous seedlings (i.e. with three fully developed leaves) were selected for each mother tree and species, corresponding to a total of 360 seedlings (60 half-sibs * 3 mother trees used as biological replicates * 2 species) and exposed to a waterlogging/drought stress cycle. An overview of the experimental design is presented in Supplementary Figure 1.

i. Waterlogging treatment: The 360 seedlings were placed in five 100 L plastic containers corresponding to five blocks. The plastic containers were filled with water that was deoxygenated directly by bubbling with N2. The seedlings were immersed in the water such that the water level reached 2 cm above the collar. The O2 concentration in each container was monitored daily with a portable O2 electrode (Cellox 325, WRW, Weilheim, Germany). Each block comprised 12 siblings from the same species and mother tree. We sampled white roots from 10 seedlings (i.e. two in each of the five blocks) for each species and each mother tree after 0 (control), 6 hours and 24 hours (short-term stress) and 9 days (long-term stress) of hypoxia. The roots were immediately frozen in liquid nitrogen and stored at -80°C until RNA extraction. The 10 seedlings were pooled for further RNA-seq analysis.
ii. Drought stress treatment: We removed the remaining 120 seedlings from the deoxygenated solution on day 10. Once the excess water had percolated through the substrate by gravity, the seedlings were maintained at field capacity for three weeks and were then subjected to drought stress (Supplementary Figure 1). Drought stress was achieved by keeping water levels in the substrate at 15% of field capacity for nine days. Before the start of the experiment, we determined the water retention capacity of the substrate by weighing the pots at field capacity and then again after drying the substrate (at 65°C for 24 hours). The amount of substrate per pot was also determined before sowing the acorns, to make it possible to measure the maximum water content per pot (∼170 g) independently of the biomass produced. Drought stress was then applied in the greenhouse, as previously described (Marguerit *et al.,* 2012). We monitored the amount of water in the substrate daily, by weighing the pots with a precision of +/- 1 g (Sartorius, Aubagne, France), until the water content in the pots reached 15% field capacity (corresponding to a weight loss of 144 g). Pots that dried faster were maintained at 15% water content for nine days before the harvesting of white roots. In parallel, we maintained a subset of seedlings at field capacity to serve as a control. We harvested white roots from these seedlings at the same time as from the stressed sample to prevent confounding ontogenetic effects (Supplementary Figure 1). As for the waterlogging treatment, we also generated three biological replicates for gene expression analysis by pooling the seedlings from the same mother tree (i.e. 10 seedlings per mother tree, Supplementary Figure 1). We also measured leaf predawn and midday water potential both for the controls and treated seedlings before each sampling, using a Scholander-type pressure chamber as described by Hsiao (1990). Measurements were made on five seedlings each for the control and the drought-stressed treatment.

### RNA extraction and sequencing

RNA was extracted and purified as described by Le Provost *et al*. (2007). Residual genomic DNA was removed with an RNase-free DNase, RQ1 (Promega, Madison, WI, USA), according to the manufacturer’s instructions, before purification. We assessed the quantity and quality of the extracted RNA on an Agilent 2100 Bioanalyzer (Agilent Technologies, Inc., Santa Clara, CA, USA).

For the waterlogging treatment, we constructed nine libraries for each species, corresponding to three biological replicates x three treatments (control, short-term response assessed by pooling equimolar amounts of total RNA extracted after six and 24 hours, and long-term response assessed on RNA extracted after nine days of waterlogging).

For the drought stress treatment, we generated six libraries for each species, corresponding to three biological replicates x two treatments (control and long-term response, i.e. after nine days of water levels at 15% field capacity in the substrate).

We generated these 30 libraries as described by Le Provost *et al*. (2016) and according to the Illumina protocol (TruSeq Stranded mRNA Sample Prep Kit). Briefly, we selected mRNA from 2 µ g of total RNA. The mRNA was then chemically fragmented and reverse transcribed with a random hexamer primer. The second strand of the cDNA was generated, 3’-adenylated, and Illumina adapters were added. We amplified DNA fragments (with adapters) by PCR with Illumina adapter-specific primers. We quantified the libraries with a Qubit Fluorometer (Life Technologies, NY, USA). We estimated their size with Agilent 2100 Bioanalyzer technology (Agilent). We then sequenced each library by 101 base- read chemistry, in a paired-end flow cell, on an Illumina HiSeq2000 (Illumina, San Diego, CA, USA). More than 24 million useable reads were generated for each library (Supplementary Table 1).

### Cleaning, mapping and identification of differentially expressed genes

We applied the following two-step procedure to identify differentially expressed genes. We first removed low-quality reads (quality value<20). The high-quality reads were then aligned with the 25,808 oak gene models previously published with the reference oak genome (Plomion et al., 2018), with the BWA mapper V0.6.1 aligner (Li & Durbin, 2009) and a maximum insert size of 600 bp, with four mismatches allowed. We then selected gene models with at least 90 reads (i.e. over all biological replicates) for the differential analysis. In the second step, we used the DESeq2 package (Love et al., 2014) to identify differentially expressed genes with a *p*-value<0.01 after adjustment for multiple testing with a false discovery rate (FDR) of 5%. The expression level of each gene was quantified by the TMM method. We considered only differentially expressed genes for which at least a two-fold change in expression was observed. For each experiment, we assessed the effects of treatment, species and their interaction in likelihood ratio tests implemented in the DESeq2 package.

The treatment and species effects were assessed by comparing a model without interaction (M1) with two simplified models for the treatment (M2) and species (M3) effects.

M1: Yijk = μ +Ti +Sj +εijk where Ti is the treatment effect (i=” control”, “short-term”, “long-term” for waterlogging and i=“control” or “long-term” for drought stress), and Sj is the species effect (j= “PO” or “SO”).

M2: Yjk = μ +Sj +εjk M3: Yik = μ +Ti +εik

For interaction effects, we compared a complete model: Yijk = μ +Ti +Sj + (S*T)ij + εijk (M4) to M1.

Differential gene expression analyses therefore yielded six genes sets: (#1) genes differentially expressed between waterlogged and control conditions (across species), (#2) genes differentially expressed between drought and control conditions (across species), (#3) genes differentially expressed between species throughout the waterlogging experiment, (#4) genes differentially expressed between species in the drought stress experiment, (#5) genes displaying significant treatment-by-species interaction in the waterlogging experiment, and (#6) gene displaying significant treatment-by-species interaction in the drought stress experiment. An additional gene set was also generated from the overlap between genes displaying a “species” effect regardless the treatment considered (gene set #7).

Annotations for the differentially expressed genes were recovered from the reference oak genome (Plomion et al.,2018).

### Geneset and subnetwork enrichment analysis

We performed GO-term enrichment analysis for the six gene sets described above with the topGO R package (Alexa, 2010). We corrected *p*-value for false discovery rate and considered ontology terms with a corrected *p*-value below 0.05 to be significantly enriched.

We then performed enrichment analysis with Pathway Studio software (Pathway Studio^®^, Elsevier 2017), as described by Le Provost et al., (2016). Our main goal was to identify genes potentially involved in ecological preferences. Therefore, in each type of experiment, we focused on the sets of genes differentially expressed between species (gene sets #3, #4 and #7) and the set of genes presenting significant treatment-by-species interactions (gene sets #5 and #6). Indeed, we expected (i) genes displaying significant treatment-by-species interactions to capture the different molecular strategies to the two water regimes evolved by each of the species, and (ii) genes displaying a species effect to have higher basal levels of expression in the tolerant species (i.e. PO for excess water and SO for water deficit). An additional network was generated with genes displaying a species effect whatever the stress considered. The previous set of genes may reveal important mechanisms underlying extrinsic barriers, whereas this last gene set should reveal genes involved in reproductive isolation (intrinsic barriers) between PO and SO.

### Reverse transcription and quantitative real-time PCR (qPCR)

For each experiment, we selected three genes displaying a significant treatment, species or interaction effect for qPCR validation. All the genes analyzed and their associated effects are listed in supplementary Table 2. We performed qPCR quantification as described by Le Provost *et al*. (2012) on 1 µ g of RNA, on a Chromo4TM Multicolor Real-Time PCR detection system (Bio-Rad Laboratories, Hercules, CA, USA) with standard PCR parameters. The fluorescence data obtained were analyzed with the Genex macro. This macro uses an algorithm derived from that described by Vandesompele et al. (2002) for qPCR normalization and quantification. We normalized fluorescence data against two control genes (Qrob_P0517760.2 encoding a hypothetical protein and Qrob_P0217660.2 encoding a ubiquitin-protein ligase RHF1A). All the primer pairs used for qPCR were designed with Primer3 software (Rozen & Skaletsky, 2000) (Supplementary Table 2). We selected control genes as follows. We first mined our digital expression analysis to identify genes that were not differentially expressed. We then selected six genes from this gene set for which coverage was similar in each biological replicate. We then evaluated the stable expression of these six genes in our biological samples with Genorm software (Vandesompele et al., 2002). We identified two genes with very stable expression levels of expression (Qrob_P0517760.2: “Unknown protein” and Qrob_P0217660.2: “Ubiquitin protein ligase RHF1A”). This validation was performed with the same trees as for RNAseq analysis, with independent RNA extractions. This analysis therefore served as a technical verification of the RNA-seq approach.

### Relative genetic divergence between differentially expressed genes

We investigated whether differentially expressed genes presented evidence of potentially adaptive genetic divergence, by assessing the overlap between the six gene sets and loci with high FST fixation indices between PO and SO. Divergence estimates were obtained with the pool-seq data recently described by Leroy et al., (2020). FST was calculated with the popoolation2 software suite (Kofler et al., 2011) for each SNP. We then assessed enrichment in high FST SNPs among the genes of each of the six gene sets. We defined high FST SNPs empirically by considering the 1, 0.5, 0.1, and 0.001% right tails of the genome-wide FST distribution. We first calculated the proportion of genes overlapping with high-FST SNPs for each gene set and each threshold. For each gene set of size N, we then built a null distribution by randomly sampling *N* genes 1,000 times, each time calculating the proportion of randomly drawn genes overlapping with high-FST SNPs. Enrichment was calculated by dividing the proportion of genes from each gene set overlapping with high-FST SNPs by the mean proportion of random genes overlapping with high-FST SNPs in the corresponding null distribution. We determined the significance of enrichment by comparing the proportion of genes from each gene set overlapping with high-FST SNPs from the corresponding null distribution.

## Results

### Characterization of the treatments applied

For waterlogging, we measured the O2 concentration in each container daily (Supplementary Figure 2A). The O2 concentration in the water in the container decreased sharply just after the start of N2 bubbling and became stable (close to 1 mg l^-1^) after 5 minutes. N2 bubbling was then stopped. The low level of O2 in the water was immediately consumed by the root system, maintaining O2 concentration in the water close to 1 mg l^-1^ throughout the experiment. The samples harvested in this experiment may therefore be considered to have been under hypoxic stress (i.e. only residual O2 in the rhizosphere). Two thirds of the seedlings (*N*=240) were harvested during the waterlogging experiment. We found visible morphological differences between the species after nine days of waterlogging stress, with hypertrophied lenticels observed only in the immersed portion of the stem of the tolerant species (Le Provost et al., 2016).

The remaining seedlings (*N*=120) were used in the drought stress experiment. We considered seedlings to be drought-stressed only if the combined weight of seedling, 0.5 L pot and soil fell to below 144 g (i.e. approx. substrate at about 15% of field capacity). We maintained all the seedlings at this threshold for nine days by weighing the pots daily and adjusting their weights with an appropriate volume of water. An analysis of the dehydration kinetics (Supplementary Figure 2B) revealed that all seedlings reached the desiccation threshold at about the same time. We measured leaf predawn and midday water potentials for each species just before sampling (Supplementary Figure 2C). We performed the same measurements on the control seedlings harvested on the same date, to prevent confounding ontogenetic effects. For both PO and SO, an increase (in absolute value) in leaf predawn and midday water potential was observed in response to stress (comparison of control and stressed samples). The midday water potentials of stressed plants were 1.5 MPa for PO and 1.7 MPa for SO. Predawn water potentials were close to 1 MPa. These results suggest that the water stress applied in this experiment was moderate, probably due to the medium used, which contained 50% peat. Bréda *et al*. (1993) reported a leaf predawn water potential value of 2 MPa for SO seedlings experiencing severe water stress. Here, we observed significant differences in leaf predawn and midday water potentials between the two treatments (i.e. control vs. stress), for both species. The difference between PO and SO under conditions of drought stress was not significant although the difference in midday water potential between the control and stressed samples was smaller for SO, suggesting a better tolerance to water stress.

### Global characterization of transcriptomic data

PO and SO seedlings were subjected to a cycle of waterlogging and drought stresses. We monitored the intensity of the constraints applied through observations and measurements throughout the experiment (see Supplementary Figure 1). Root tips were then sampled and 30 cDNA libraries were constructed from mRNA extracted from these tissues: 18 libraries for waterlogging and 12 for drought conditions. For each library, 23 to 43 million reads were generated. An overview of the sequenced libraries is shown in Supplementary Table 1, along with their SRA accession numbers For the waterlogging experiment, three sampling points were analyzed (control C, short term ST, and long term LT responses) for both species. For the drought stress experiment, we analyzed two sampling points in both species (control and long-term response, after 9 days of water deficit). We obtained a mean Phred score of 35 for the libraries analyzed in this study. Finally, we successfully mapped 82% to 88% of the sequences from each library onto oak gene models.

We investigated the reproducibility of biological replicates and the correlation between the various samples by performing principal component analysis (PCA) on the waterlogging (18 samples, Figure 1A) and drought stress (12 samples, Figure 1B) datasets. This descriptive analysis showed close clustering within biological replicates, attesting to the quality of the RNAseq data. It also showed that the samples (species x treatment) clustered into different groups corresponding to the different combinations analyzed in this study. However, whereas the PCA for the drought stress samples displayed clear differences in expression between species and between treatments (Figure 1B), that for the waterlogged samples revealed clear differences only between the three treatments (Figure 1A). Subtle differences in terms of the number of genes and/or changes in expression between species therefore probably remain to be discovered by other approaches.

**Figure 1:**
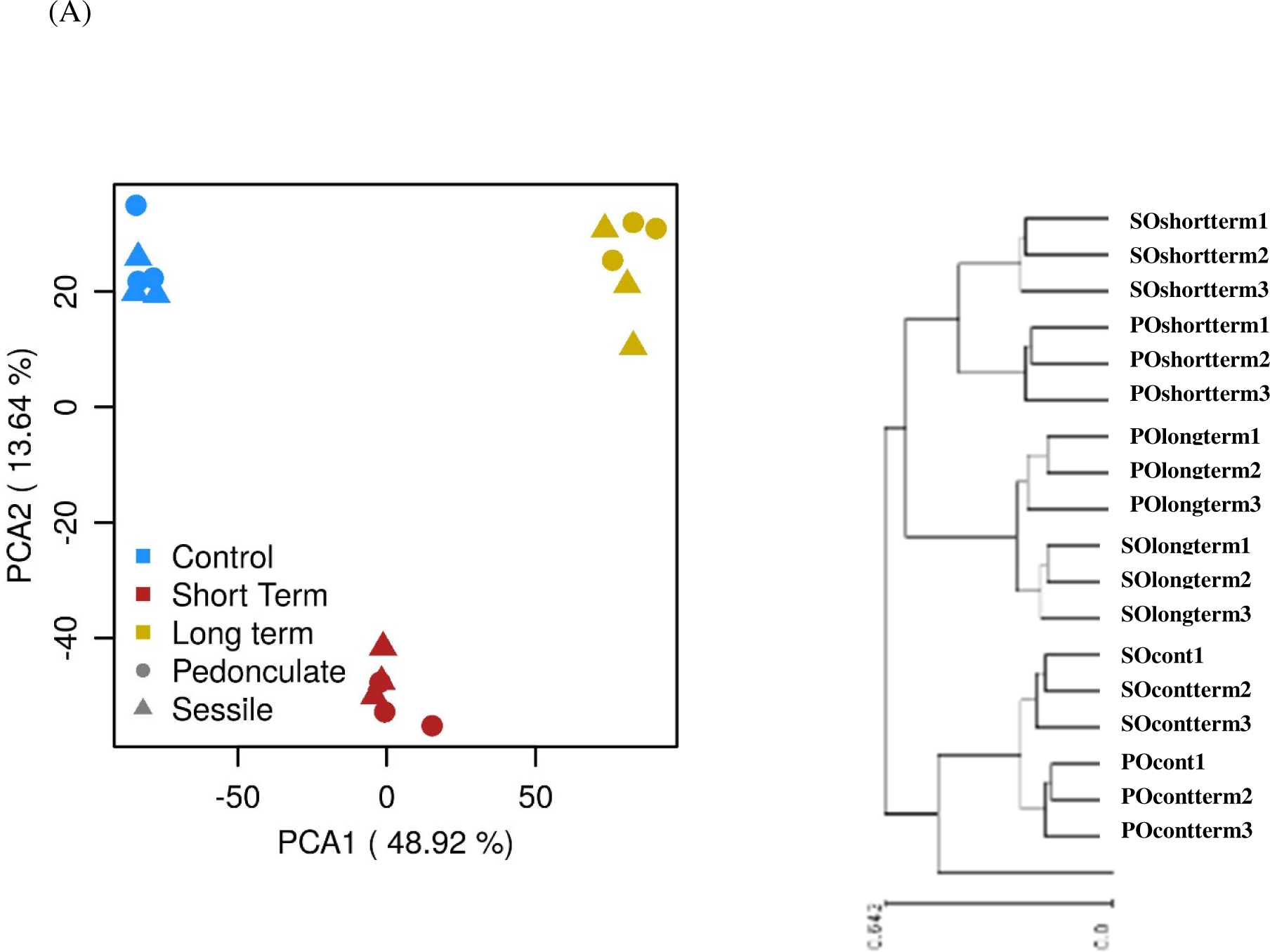
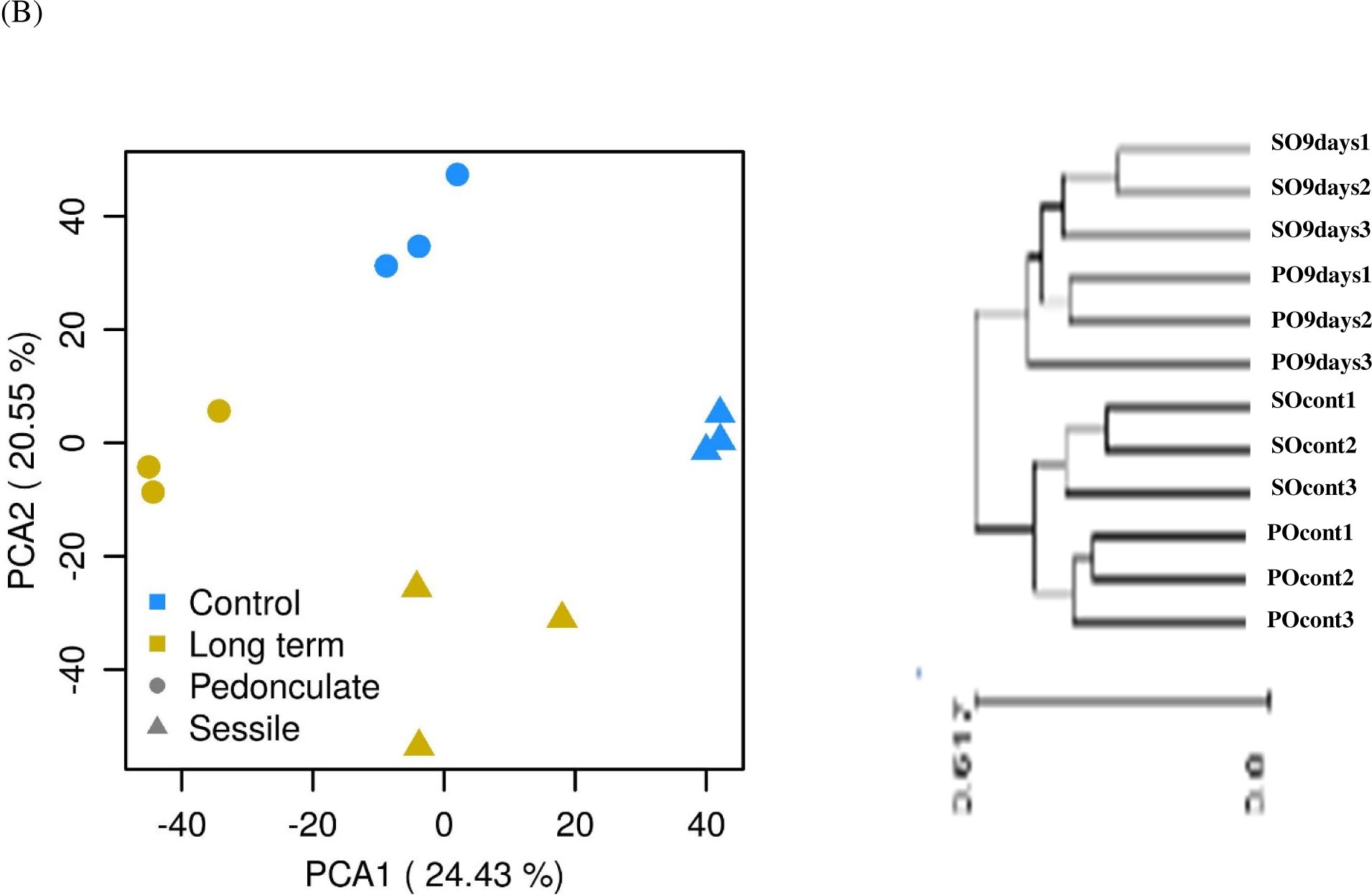
PCA and hierarchical clustering analysis.

The adjusted data were also evaluated with a clustering approach in Expender software (Shamir et al., 2005). This analysis confirmed the high degree of reproducibility for the RNAseq data, with biological replicates grouped together in the same cluster (Figure 1).

### Number and effect size of differentially expressed genes

DEGs (i.e. with an absolute fold-change in expression ≥2 and adjusted *p*-values ≤0.01) were identified separately for the waterlogging and drought stress experiments with the DESeq2 package. We used the likelihood ratio test to identify genes displaying significant “species”, “treatment” and/or “interaction” effects.

For the waterlogging experiment, 18,155 genes covered by more than 90 reads remained for abundance analysis. Most of these genes (18,146, 99.9%) were covered by sequences from both species. We identified five genes specific to PO (i.e. covered by PO reads only): Qrob_P0335670.2 (germacrene-D-synthase), Qrob_P0495260.2 (glucose-induced degradation protein), Qrob_P0281130.2 (kinase with adenine nucleotide alpha hydrolase-like domain), Qrob_P0336470.2 (PPPDE thiol peptidase family protein), Qrob_P0125380.2 (SABATH methyltransferase). We also identified four genes specific to SO: Qrob_P0362510.2 (myb-like HTH transcriptional regulator family protein), Qrob_P0119020.2 (cysteine-rich repeat secretory protein 38), Qrob_P0598470.2 (subtilase family protein T6P5), and Qrob_P0605080.2 (14-3-3-like protein GF14 omega protein).

We also identified 820 genes with a significant treatment effect, 1,167 genes with a significant species effect, and 319 genes with a significant interaction (Table 1, Supplementary Files 1, 2, and 3 and Figure 2A and 2B). Only 25 genes had all three types of effect (Figure 2B), but 183 genes had a significant treatment effect only, 522 had a significant species effect only and 278 genes had an interaction effect.

**Figure 2:**
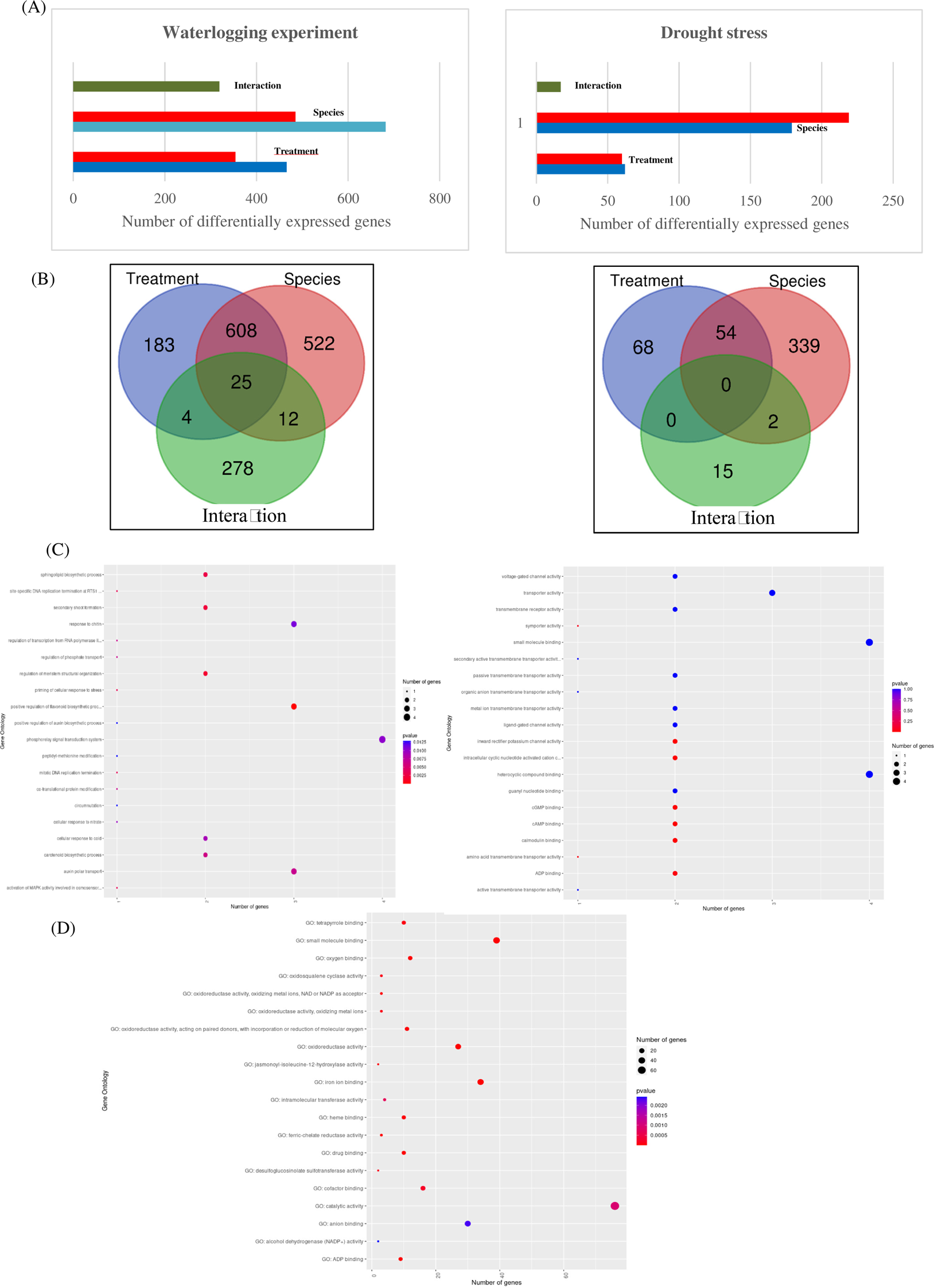
Illustration of the main results obtained from the DEGs analysis. Panel A: Bar charts illustrating the number of DEGS obtained for each effect both for waterlogging (left section) and drought stress (right section). Panel B: Venn diagram of the genes showing a significant treatment, species or interaction effect. Left panel: waterlogging experiment, Right panel: Drought stress experiment. Panel C: Most enriched ontologies (BP) obtained for genes displaying a significant treatment-by-species interaction both for waterlogging (left part) and drought stress (right part). Panel (D) Most enriched ontologies (BP) obtained for genes displaying a significant species whatever the stress analyzed.

**Table 1:** Summary of the DESeq2 analysis for the two experiments (waterlogging and drought). Abbreviations: PO: pedunculate oak, SO: sessile oak. The number of genes in each cell was determined with the following parameters: adjusted *P* value < 0.01 and fold change > or = 2. For each effect, the compared models *Mi* (see materials and method section) are indicated in parentheses.

|  | WATERLOGGING |  |  |
| --- | --- | --- | --- |
|  | Treatment (M1-M2) | Species (M1-M3) | Interaction (M4) |
| PO | 466 | 682 | NA |
| SO | 354 | 485 | NA |
| **TOTAL** | **820** | **1,167** | **319** |
|  | Drought |  |  |
|  | Treatment (M1-M2) | Species (M1-M3) | Interaction (M4) |
| PO | **62** | **179** | **NA** |
| SO | **60** | **216** | **NA** |
| **TOTAL** | **122** | **395** | **17** |

For the drought stress experiment, 17,688 genes were considered according to the criteria defined above. We identified 11 genes specific to either PO or SO: two genes for PO (Qrob_P0374520.2 (unknown protein) and Qrob_P0084850.2 (cylic nucleotide gated channel 1) and nine genes for SO (Qrob_P0711010.2 (unknown protein), Qrob_P0675500.2 (aspartyl protease family protein), Qrob_P0092630.2 (TIR-NBS-LRR class disease resistance protein), Qrob_P0627090.2 (HXXXD-type acyl-transferase family protein), Qrob_P0446600.2 (NDR1/HIN1-like protein), Qrob_P0169890.2 (unknown protein), Qrob_P0684400.2 (cycloartenol synthase), Qrob_P0362510.2 (myb-like HTH transcriptional regulator family protein) and Qrob_P0593920.2 (DUF946 family protein).

Application of the same statistical criteria as for the waterlogging experiment led to the identification of 122 genes with a significant treatment effect, 395 genes with a significant species effect, and 17 genes with a significant interaction (Table 1, Supplementary File 4, 5 and 6, Figure 2A and 2B). The overlap between these three types of effects is shown in the Venn diagram of Figure 2B. No gene displayed all three types of effect simultaneously, and we found 68 genes with a significant treatment effect only, 339 genes with a significant species effect only and 15 genes with a significant interaction.

We found that 184 of the genes displaying a “species” effect (i.e. 13.4% of the “species” effect genes) were differentially expressed in both experiments (Supplementary File 7). This overlap is highly significant overlap (chi² test on a 2x2 contingency table: chi² = 1154, *p*-value < 2.2e-16). These 184 genes displaying a species effect whatever the stress considered are potentially relevant as genes involved in reproductive isolation (intrinsic barriers) between PO and SO. These genes are listed in supplementary file 7.

### Validation of RNA-seq results by qPCR

We performed reverse transcription-quantitative PCR analysis for 10 genes (six identified in the waterlogging experiment and four identified in the drought stress experiment, respectively). The expression patterns of the tested genes revealed by qPCR were similar to those obtained by RNAseq (see supplemental information 1), confirming the reliability of the RNA-seq data.

### Gene set and subnetwork enrichment analyses

The main goal of this study was to highlight the molecular mechanisms involved in adaptation to waterlogging or drought stress, in PO and SO, respectively, rather than to identify genes showing a plastic response to these constraints. We therefore chose to perform gene set and subnetwork enrichment analyses exclusively on the genes displaying a significant interaction (in either experiment, i.e. gene sets #5 and |#6) or species (across experiments, i.e. gene set #7) effect. Indeed, we hypothesized that the interaction term would reveal genes defining species-specific adaptive molecular strategies underlying reproductive barriers, whereas genes displaying a “species” effect, whatever the treatment considered, would reveal intrinsic functions driving the reproductive isolation between PO and SO, regardless of environmental conditions. Genes displaying a “species” effect in specific environmental conditions (gene sets #3 and #4) may also constitute an important source of information because higher basal levels of expression in the tolerant species may explain its better adaptation to the environmental constraints with which it is usually confronted in natural conditions. Below, we focus exclusively on the set of genes displaying a “species” effect across experiments. The species-specific genes displaying a response to either waterlogging or drought stress separately are presented in Supplemental Information 2.

#### (i)#Gene set enrichment analysis

The results are shown only for the first 100 gene ontologies in Supplementary files 3, 6 and 7. A graphical representation of the first 20 ontologies is also available in the same supplementary Files for a clearer view of the ontologies regulated.

For interaction-responsive genes, the highest levels of enrichment during waterlogging were observed for the biological processes (BPs) “positive regulation of flavonoid biosynthetic process”, “regulation of meristem structural organization”, “secondary shoot formation”, “sphingolipid biosynthetic process” and “activation of MAPK activity involved in osmosensor”, and for the molecular functions (MFs) “phosphorelay response regulator activity”, “ubiquitin-protein transferase activity”, “peptide deformylase activity”, “sphingolipid delta-8 desaturase activity”. Very different BPs (“regulation of membrane potential”, “calcium ion transport”, “potassium ion transport” and “amino acid import”) and MFs (intracellular cyclic nucleotide activated cation channel”, “inward rectifier potassium channel activity”, “cAMP binding”, “cGMP binding” and “calmodulin binding”) were found to be enriched during drought conditions (Supplementary File 3 and 6 and Figure 2C).

The results of the gene set enrichment analysis for genes displaying a “species” effect whatever the treatment considered are shown in supplementary file 7 and Figure 2D. The BPs “oxidation-reduction process”, “response to stimulus”, “defense response”, “triterpenoid biosynthetic process” and “triterpenoid metabolic process” were found to be enriched, and enrichment was observed for the MFs “oxygen binding”, “ADP binding”, “oxidoreductase activity”, “iron ion binding” and “drug binding”.

(ii) Subnetwork enrichment analysis was performed to highlight key molecular players involved in adaptation to waterlogging in PO or drought stress in SO. A network based on interaction-responsive genes was generated for each treatment. The functional networks are shown in Figure 3 for waterlogging and Figure 4 for drought stress. The genes with an interaction effect in waterlogged conditions formed several important hubs related to “root growth”, “response to auxin stimulus”, “jasmonate response”, “meristem function”, “cell expansion”, “plant defense” and “cell death”. Different hubs were identified for drought stress, with molecular functions related to “defense response”, “response to auxin stimulus”, “growth yield” and “seed yield”.

**Figure 3:**
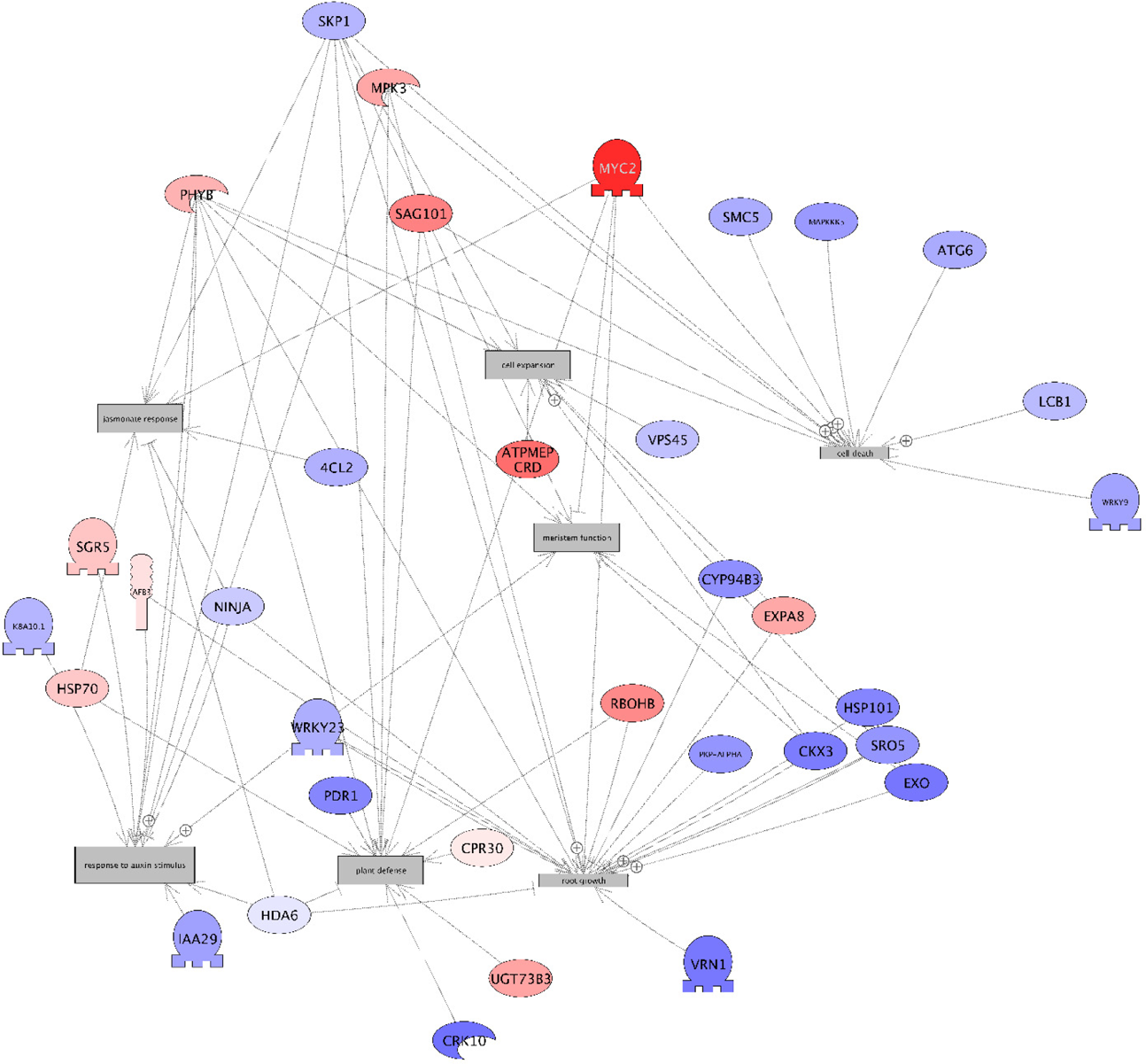
Functional network for genes displayed a significant treatment-by-interaction effect in the waterlogging experiment.

**Figure 4:**
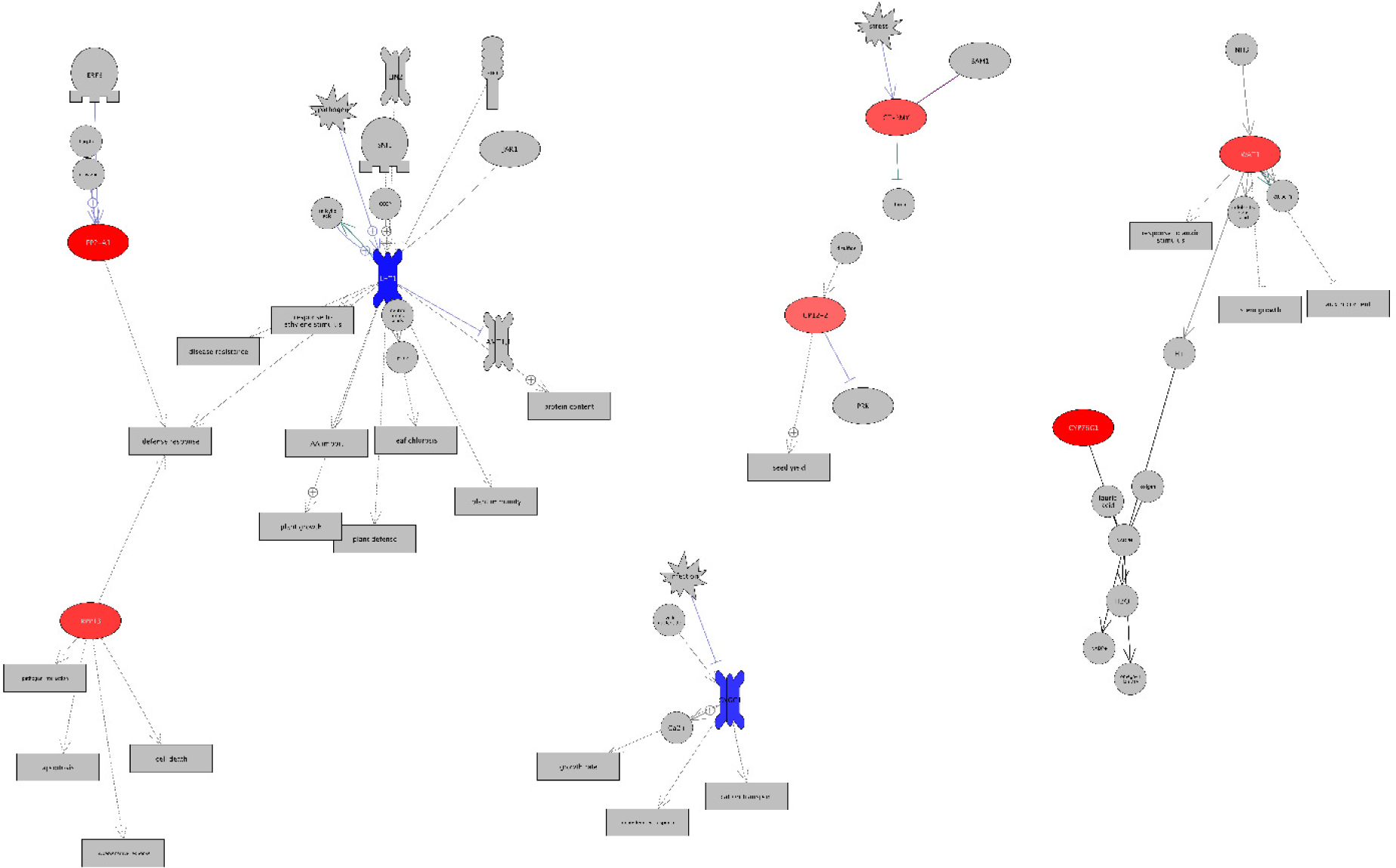
Functional network for genes displayed a significant treatment-by-interaction effect in the Drought stress experiment.

An additional subnetwork was also generated from the genes displaying a “species” effect whatever the treatment analyzed. This network is shown is Figure 5. We identified hubs related to: “flowering time”, “cell differentiation”, “cell proliferation”, “developmental process” and “seed germination”. Meristematic cells generally display an enrichment in these molecular functions. This result adds weight to the hypothesis that this last gene set contains genes potentially involved in the formation of intrinsic barriers driving the reproductive isolation between PO and SO, regardless of environmental conditions.

**Figure 5:**
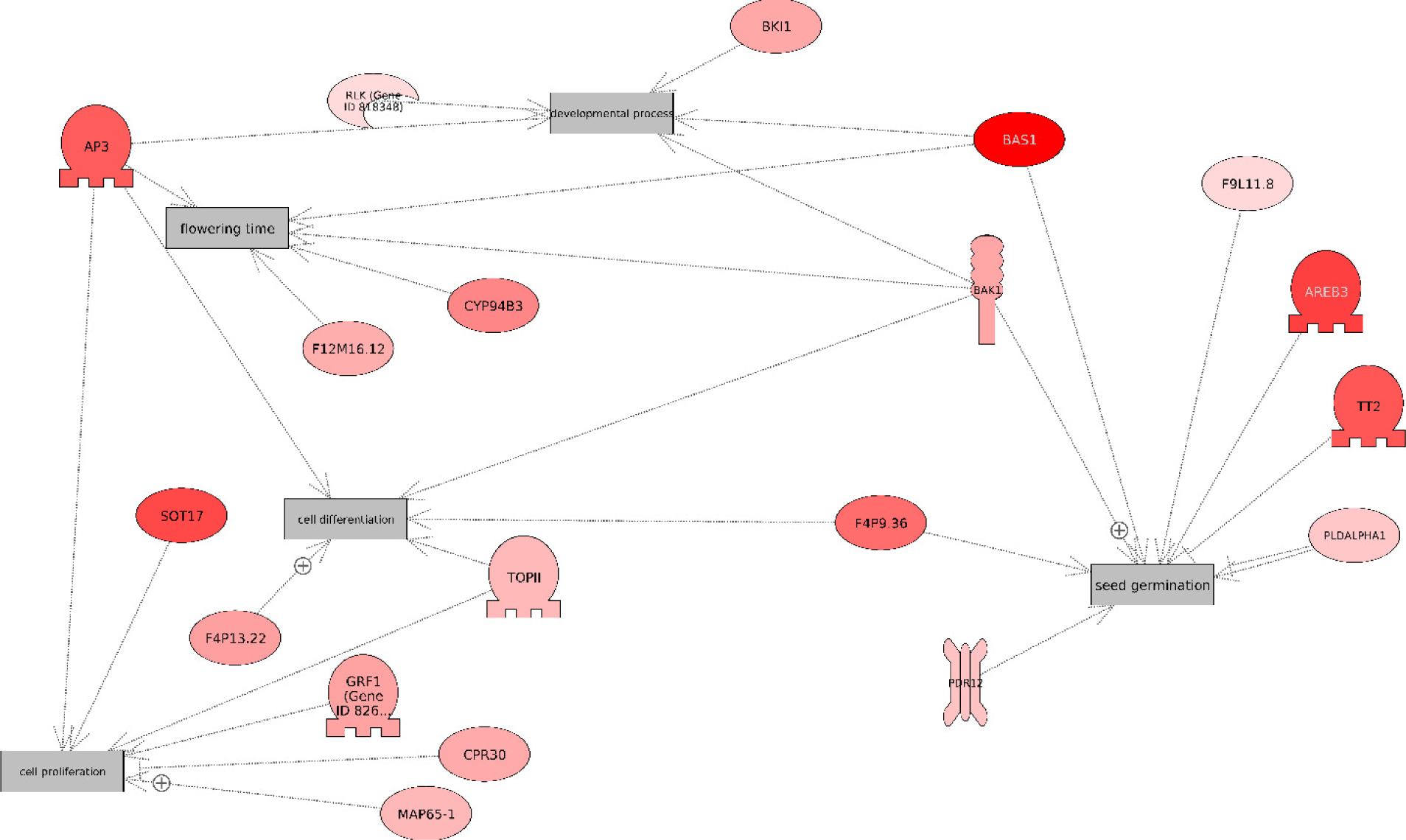
Functional network for genes displayed a significant species effect whatever the stress applied.

### Overlap between loci showing a high degree of genetic differentiation between species and differentially expressed genes

For this comparison, we used a published dataset from a study investigating adaptive differentiation between PO and SO based on pool sequencing (Leroy et al., 2020). The mean FST value across the 39,644,639 SNPs with FST>0 was 0.0545, suggesting weak overall genetic differentiation. However, FST estimates ranged from 1.17e^-6^ to 1. We assessed enrichment in gene sets identified by differential expression analysis for SNPs in the 1, 0.5, 0.1, 0.01, and 0.001% right-hand tails of the genome-wide FST distribution, corresponding to thresholds of 0.14, 0.217, 0.454, 0.803, and 1, respectively (Figure 5). In each of the six gene sets (#1 to #6), we found significant enrichment in SNPs displaying a high degree of differentiation between the two oak species (Figure 6). Enrichment ratios exceeded six-fold for the species and treatment effects in the drought experiment, but all six gene sets displayed significant enrichment, suggesting that the sets of genes differentially expressed in our experiments included genes related to adaptive differentiation between the two species. The lists of genes including highly differentiated markers for the various thresholds are provided in Supplementary File 8.

**Figure 6:**
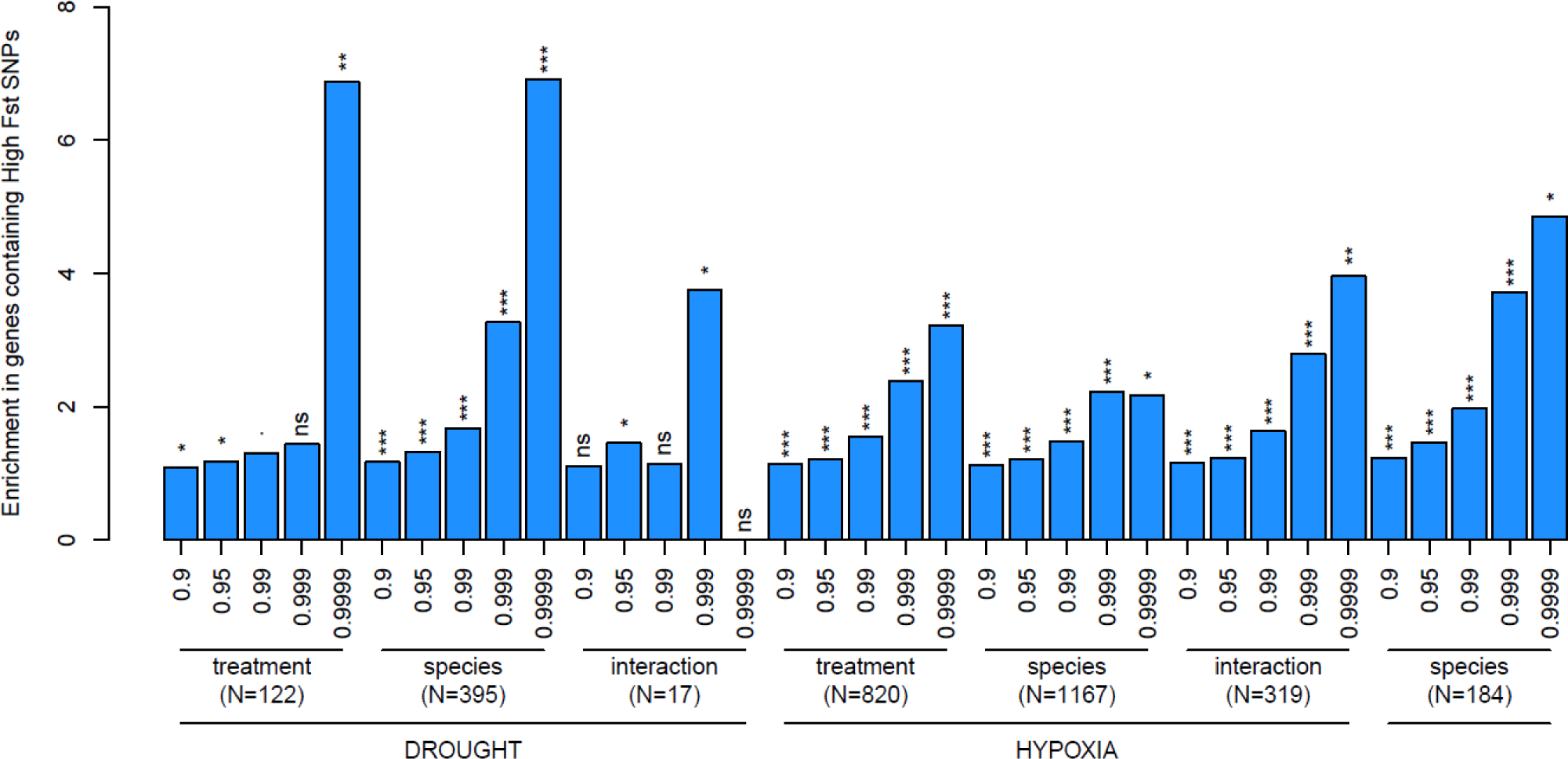
Genes differentially expressed between treatments and species in the drought and water- logging experiments are enriched in highly differentiated single nucleotide polymorphisms (SNPs) between *Q. robur* and *Q. petraea*. SNPs differentiation between the two species *Q. robur* and *Q. petraea* were obtained from Leroy et al., (2019). The y-axis represents the enrichment of the seven differentially expressed genes sets investigated in this study (x-axis). For example, an enrichment of above six for genes differencially expressed between treatments in the Drought experiments indicates that this gene set included over six times more genes with highly differentiated SNPs than expected by chance. For each genes set, the five bars correspond to enrichment for genes including SNPs with FST values above the 0.9, 0.95, 0.99, 0.999, 0.9999 quantile of the genome-wide FST distribution, respectively. Significance of enrichment estimates were assessed using 1000 permutations and are indicated above each bars: ns: non-significant, .: p-value < 0.1, *: p-value <0.05, **: p-value <0.01, ***: p-value <0.001.

We also assessed the enrichment in high-FST SNPs among genes displaying differential expression between species and across treatments (Figure 6, rightmost barplot). Again, we found a significant overlap, suggesting potential involvement in intrinsic barriers to reproduction between PO and SO.

## Discussion

The few studies published to date on the molecular plasticity of European white oaks during waterlogging or drought stress were all performed in singles species (Folzer et al., 2006; Parent et al., 2008; Spieß et al., 2012), limiting the significance of the results produced with respect to the role of gene expression in species ecological preferences. Moreover, these studies were generally performed with targeted genomic approaches and therefore focused on a limited number of genes, again reducing the scope of the findings concerning the molecular mechanisms driving oak adaptation. We investigated the contribution of gene expression to the divergence of PO and SO in a factorial design in which each species was subjected to two sets of water stress (excess or deficit) corresponding to the environmental constraints to which these two species are better adapted. This made it possible to focus on the “species x treatment” interaction term and to reveal species-specific adaptive molecular features presumably involved in the ecological preferences of these species (319 DEG for waterlogging and 17 for drought stress). This experiment also made it possible to identify genes expressed differentially between species regardless of the treatment imposed (184 genes), suggesting intrinsic roles in driving the reproductive isolation between PO and SO. In addition to these two sets of DEGs (the focus of the discussion), genes differentially expressed between the species in each treatment may also provide an important source of information, because genes with higher basal levels of expression in tolerant than in intolerant species may contribute to adaptation to a specific environmental condition. A dedicated discussion on these genes is provided in Supplemental Information 2.

### Molecular mechanisms underlying the ecological preferences of PO and SO

We focus here on the interaction-responsive genes identified in the waterlogging and drought stress experiments. We hypothesized that genes with expression levels significantly affected by the species x treatment interaction would include genes involved in the respective ecological preferences of the two species that could potentially be involved in ecological barriers.

#### Waterlogging experiment

Waterlogging had a stronger effect than drought stress in terms of the number of DEGs (319) displaying “species x treatment” interactions. The subnetwork generated (Figure 3) revealed important hubs related to root growth (“root growth”), hormone signaling (“response to auxin stimulus” and “jasmonate response”), cell differentiation and expansion (“meristem function” and “cell expansion”), plant defense and cell death, suggesting a key role for molecular mechanisms triggering the formation of aerenchyma and adventitious root formation. In PO, Parelle et al., (2006) reported that morphological and anatomical changes (such as the formation of adventitious roots and aerenchyma) were essential to cope with hypoxia. They also reported the formation of larger amounts of these adaptive structures in PO than in SO. This formation of aerenchyma or adventitious roots in waterlogging-tolerant plants may be triggered by genes belonging to the main hubs identified here, suggesting that the molecular mechanisms identified here are of particular importance for the adaptation of PO to waterlogging. We review some of the genes identified below. In the following section, we focus our discussion on some genes taking into account their Fst value.

##### (1)#Programmed cell death leading to aerenchyma formation

Programmed cell death (PCD) is a major mechanism underlying plant development and adaptation to abiotic or biotic stress (Smith et al., 2015). In particular, it is involved in the modification of root architecture under drought stress and the formation of aerenchyma in hypoxic conditions. Tolerant plants typically produce aerenchyma in response to hypoxic conditions through the lysis and death of cells in the root cortex to produce additional gas space (Yamauchi et al., 2013). The tolerance to waterlogging observed in PO may be partially explained by its ability to produce more aerenchyma than SO (Parelle et al., 2006), a finding confirmed by our network analysis, which identified several genes related to cell death, a process essential to aerenchyma formation (Figure 4). The genes upregulated in PO included (i) *LCB1* (*Qrob_P0060010.2, FST:0.43*) encoding a long-chain base 1 (LCB1) subunit of serine palmitoyltransferase. In *Arabidopsis thaliana,* this gene has been shown to regulate PCD by inducing reactive oxygen intermediates (Shi et al., 2007), (ii) ATG6 (*Qrob_P0479390.2, FST:0.19*) which encodes a core autophagy-related protein involved in the regulation of autophagy (Xu et al., 2017), (iii) WRKY 9 (Qrob_P0436950.2, FST: 0.24), which is known to be involved in cell death regulation in tobacco (Liu and He, 2017) and (iv) SKP1 (Qrob_P0747290.2, Fst:0.2) encoding S-phase kinase-associated protein 1 and including multiple differentiated SNPs with a maximum FST value of 0.73. To our knowledge, this is the first report linking potentially SKP1 protein to aerenchyma formation.

##### (2)#Root growth and phytohormone biosynthesis

The formation of adventitious roots is another major morphological response to waterlogging in tolerant species. These roots contain aerenchyma (i.e. air channels), which favors the diffusion of gases in conditions of submersion (Evans, 2003). We identified important hubs potentially involved in the establishment of adventitious roots (root growth, cell expansion, meristem function) (Figure 3). We also detected hubs related to phytohormones (response to auxin stimulus and jasmonate response) known to play a key role in the formation of adventitious roots. Our analysis thus highlighted molecular networks related to this morphological adaptation in PO.

Our analysis highlighted genes (upregulated in PO) involved in either primary root growth inhibition or lateral root development. A first subset of these genes is involved in auxin signaling *(HDA6*, *WRKY23*, *AFB3*) or jasmonate responses (*PHYB*, *SKP1*, *4CL2*, *MYC2*) consistent with the modification of the root system during hypoxic responses in PO being driven by phytohormones. For instance, *MYC2* (*Qrob_P0056680.2,* FST: 0.24) encodes a basic helix-loop-helix (bHLH) DNA-binding protein. Kazan and Manners (2013) identified this gene as a master regulator of many aspects of jasmonate signaling playing a strong role in the formation of lateral and adventitious roots. *EXO* (Qrob_P0764920.2, FST:0.14) is under the control of brassinosteroid. Its overexpression in *Arabidopsis* has been reported to promote shoot and root growth (Schröder et al., (2009)).

*RBOHB* (*Qrob_P0493470.2,* FST: 0.28), which encodes a respiratory burst oxidase protein involved in the production of reactive oxygen species, is also upregulated in PO. The reactive oxygen species (ROS) produced by the product of *RBOHB* genes are involved in several developmental processes. Montiel et al., (2013) reported that the overexpression of this gene promotes lateral root growth, suggesting a potential role in adventitious root formation in PO. We also identified *NINJA* (*Qrob_P0009970.2,* FST: 0.72) as being more strongly expressed in PO. This gene encodes an interactor of the jasmonate Zim domain 10. In a study of *Arabidopsis* mutants, Acosta et al. (2013) reported a downregulation of jasmonate signaling by NINJA that was associated with a large decrease in the size of the root system. We hypothesize that the overexpression of *NINJA* in PO is associated with its ability to maintain adventitious root formation.

Two other genes were found to be downregulated in PO: *HDA6* (*Qrob_P0451700.2,* FST: 0.82), which encodes an RPD3-type histone deacetylase, a key enzyme involved in the jasmonate response (Wu et al., (2008), and *CYP94B3* (*Qrob_P0369790.2,* FST: 0.49), which encodes a cytochrome P450 protein involved in jasmonate catabolism. Heitz et al. (2012) reported a correlation between *CYP94B3* overexpression and insensitivity to jasmonate and a smaller lateral root system. These results highlight a stronger inhibition of the jasmonate response in PO, which is essential for lateral root formation, potentially accounting for the increase in lateral root system size in PO during waterlogging.

##### (3)#Meristem function and response to auxin stimulation

The *WRKY23* gene (*Qrob_P0697030.2,* FST: 0.59), which is involved in the auxin response, was more strongly expressed in PO. Grunewald et al., (2012) reported an important role for this gene in the establishment of an auxin gradient in the root system, allowing the maintenance of root meristematic activity and lateral root formation.

The *AFB3* gene (*Qrob_P0036220.2, F*ST*: 0.75*), which encodes an auxin receptor F box protein, was also upregulated in PO. *AFB* genes belong to a multigene family regulated by auxin and involved in several plant developmental processes. In *Arabidopsis thaliana,* Vidal et al. (2010) showed that the *AFB3* mutant was characterized by altered lateral root growth, whereas the primary root system was unaffected, suggesting a key role for AFB3 in modulating the architecture of the root system.

Overall, our network analysis highlighted several molecular mechanisms potentially involved in the formation of adaptive structures known to favor oxygen diffusion in the root system (aerenchyma and adventitious roots). This suggests that molecular mechanisms analogous to those observed in other tolerant plant species are upregulated in PO relative to SO.

### Drought stress

The functional network obtained from the 17 genes found to be differentially expressed in response to drought is presented in Figure 4. Three genes upregulated in SO were considered to be of particular importance (*PP2-A1*, *CT-BMY*, and *WAT1*). The *WAT1* gene (*Qrob_P0331600.2,* FST: 0.23) encodes a vacuolar auxin transporter. Auxin plays a crucial role in abiotic stress responses controlling vascular development (Ranocha et al., 2013). Zhang et al. (2017) reported that this gene was upregulated in drought-tolerant maize lines during drought stress. We also identified a *CT-BMY* gene (*Qrob_P0567500.2*, FST: 0.25) encoding a beta-amylase known to be upregulated during abiotic stress. The upregulation of this gene is generally correlated with maltose accumulation, which protects proteins, membranes and the photosynthetic electron transport chain during water stress (Kaplan & Guy, 2004). Finally, we identified *PP2-A1* (*Qrob_P0654080.2,* FST: 0.36), a gene involved in the plant defense response. Zhang et al. (2011) reported a role for PP2-A1 in the activation of plant defense responses during biotic stress.

In conclusion, our functional network analysis identified drought-responsive genes in SO related to three major pathways: ABA response, osmoregulation and plant defense, all of which are known to be upregulated in drought-tolerant genotypes. These results suggest that SO has a better capacity to maintain both its primary root growth and osmoregulation, facilitating the maintenance of water metabolism during water shortage and greater tolerance to drought stress. In addition, most of the genes highlighted by our functional network presented at least one SNP above the 90% quantile of the genome-wide FST distribution, suggesting that our functional analysis yielded genes not only important for the response to stress in SO, but also probably subject to strong positive selection.

### Molecular mechanisms underlying intrinsic reproductive isolation barriers

We focus here on the genes displaying a “species” effect regardless of the treatment imposed. We hypothesize that this gene set contains key molecular players potentially involved in intrinsic barriers to reproductive isolation. Indeed, our functional analysis (Figure 5) highlighted several processes (i.e. “flowering time”, “cell proliferation”, “cell differentiation”) related to this biological function. Our transcriptome data were obtained from roots rather than floral meristems, but the molecular mechanisms identified here probably reflect important functions of plant meristems generally. This hypothesis is also supported by the high level of differentiation of the genes identified in our biological network, ranging from 0.15 (F9L11.8) to 1 (BAS1). Moreover, 164 of the 184 genes identified in the whole dataset (89%) were also found in the 0.1% tail of the genome- wide FST distribution, with max Fst values ranging from 0.14 to 1 (Supplementary File 8). Some of these genes may, therefore, correspond to intrinsic reproductive barrier loci.

Eight genes in the biological network shown in Figure 4 were considered to be of particular interest. Four of these genes relate to the “flowering time” process: *BAS1* (Qrob_P0551020.2, Fst: 1), *AP3* (Qrob_P0454040.2, Fst: 0.64), CYP94B3 (Qrob_P0369770.2, Fst: 0.47) and *F12M16.14* (Qrob_P0657620.2, Fst: 0.31). Sandhu et al. (2012) showed that the *BAS1* gene encodes a cytochrome P450 protein involved in brassinosteroid catabolism and reported a key role for brassinosteroid inactivation during the floral induction of AP3, a key component of meristem identity in plants. AP3 interacts with AP1 to specify floral meristem identity during floral transition in *Arabidopsis thaliana* (Liu et al., 2007). CYP94B3 also encodes a cytochrome P450 protein. Bruckhoff et al., (2016) suggested that this gene might play an important role in controlling flowering time by inactivating the phytohormone jasmonyl isoleucine. We also identified two other CYP proteins with high Fst values in our gene list (but not in our functional network): CYP72A9 (Qrob_P0084340.2, Fst: 1) and CYP76G1 (Qrob_P0088450.2, Fst: 0.75). These findings suggest that some members of this group may be major targets of natural selection. Finally, we identified an F12M16.14 gene encoding a Na^+^/Ca ^2+^ exchanger-like protein involved in maintaining Ca^2+^ homeostasis. Li et al., (2016) reported that transgenic *Arabidopsis* lines displayed alterations to flowering time due to changes in the expression of two major flowering genes (the *Constans* and *Flowering* loci).

Three of the genes identified in our network analysis (*AREB3* (Qrob_P0187720.2, Fst: 0.75) *TT2* (Qrob_P0304680.2, Fst: 0.66) and *F4P9.36* (Qrob_P0554070.2, Fst: 0.57)) belong to the “seed germination process” (Figure 5): i) *AREB3* encodes an ABA-responsive element binding protein with a bZIP domain. Wang et al. (2015) reported a key role for this gene in germination, seed development and embryo maturation. Moreover, Hoth et al. (2010) showed that *AREB3* is co-expressed with a sugar transporter gene (*AtSUC1)* in pollen, ii) *TT2,* encoding an R2R3 Myb transcription factor involved in anthocyanin/proanthocyanin accumulation was also identified. Zhao et al. (2019) reported that *Arabidopsis* mutants with a downregulated TT2 gene had a shorter dormancy period, suggesting an important role for this gene in meristem functioning, and iii) *F4P9.36*, encoding a protein similar to dihydroflavonol-4-reductase. Østergaard et al. (2001) showed that this protein was involved in embryogenesis and seed maturation in *Arabidopsis thaliana*.

Finally, we identified a SOT17 gene (Qrob_P0168490.2, Fst: 0.72) associated with cell proliferation. SOT17 is similar to desulfoglucosinolate sulfotransferase. The SOT17 gene has been reported to be expressed in the shoot apical meristem and is potentially involved in cell proliferation in the root meristem via the glucose TOR signaling pathway (Kim et al., (2017); Xiong & Sheen (2013)).

However, most of these 184 genes were not found in our subnetwork enrichment analysis (166 of 185). Below, we focus exclusively on those with high Fst values.

The genes with high Fst values included three genes (two CAS1 genes (Qrob_P0684390.2, Fst:1) and a NOT2 gene (Qrob_P0689750.2,Fst 0.77)) involved in male gametophyte initiation. Wang et al., (2013) reported that the downregulation of *NOT2* in *Arabidopsis* caused severe defects in the male gametophyte. We also identified two CAS1 gene encoding a cycloartenol synthase 1 involved in sterol biosynthesis. Babiychuk et al., (2008) showed that the CAS1 gene have a key role in the male gametophyte function and in the meristematic activity. We identified a F14M4.3 gene (Fst: 0.92) similar to a short-chain dehydrogenase reductase 5 (SDR). Cheng et al., (2002) reported that the SDR super family is involved in both hormone biosynthesis in mammals and sex determination in maize. Three BGLU17 genes with Fst values ranging from 0.97 to 1 (Qrob_P0491340.2, Qrob_P0491360.2 and Qrob_P0597130.2) were also found in our analysis. BGLU17 encodes beta glucosidase proteins potentially involved in meristem functioning through the initiation of cell division. Brzobohaty et al. (1993) reported that beta glucosidase cleaved the cytokinin-O-glucoside conjugates, which are biologically inactive, to release active cytokinin, which is essential to initiate cell division in the meristem. A PAP2 gene (Qrob_P0010280.2, Fst: 0.79) encoding a protein highly similar to phytochrome-associated protein 2 was also found in our gene list. Kobayashi et al. (2010) showed that the pattern of meristem initiation was disorganized in rice mutants for this gene. In the mutant transgenic lines, new meristems were unable to develop into spikelet meristems, highlighting a key role for PAP2 in the initiation of floral meristems. Finally, an F13K3.9 (Qrob_P0012240.2, Fst: 0.89) gene encoding a 2-oxoglutarate (2OG) and Fe(II)- dependent oxygenase protein was identified. Ciannamea et al. (2006) showed that this gene was downregulated during vernalization, suggesting a possible role in floral meristem initiation.

In conclusion, our analysis highlighted several genes potentially involved in flowering time regulation, meristem functioning, embryo maturation, cell proliferation, male gametophyte initiation and floral meristem identity. All the genes discussed in this section had high Fst values, suggesting that they are potential targets of natural selection. Overall, these results provide support for a potential role of these genes in maintaining intrinsic barriers to reproduction between oak species.

## Supporting information

Validation of RNA-seq results by qPCR.

Species responsive genes in the waterlogging and drought stress experiments.

Ecophysiological characterization of the experimental design. Evolution of the O2 concentration, water content in the substrate and water potential du

Overview of the cDNA libraries constructed in this study.

List of the primer pairs used for qPCR analysis.

Schematic representation of the experimental design used in this study.

## Acknowledgments

This work was supported by INRAE, the Genoscope: the *Commissariat à l’Energie Atomique et aux Energies Alternatives* (CEA) and ANR (GENOAK project 2011 BSV6 009 01). We thank the Genotoul Bioinformatics Platform Toulouse Occitanie (Bioinfo Genotoul, https://doi.org/10.15454/1.5572369328961167E12) for providing computing resources, and the Genome Transcriptome Facility of Bordeaux (grant from I*nvestissements d’Avenir, Convention attributive d’aide* EquipEx Xyloforest ANR-10-EQPX-16-01) for providing the infrastructure for RT-qPCR. IL received funding from INRAE and TB received funding from the ANR and the European Union’s *ERC program (*TREEPEACE # FP7-339728).

## Data accessibility

All sequences generated in this study were deposited in the Short Read archive of NCBI under PRJEB17875 (Waterlogging experiment) and PRJED19536 (Drought stress experiment) accession numbers.

## Author contributions

GLP and CP designed the study. GLP, IL, BB and CP wrote the manuscript. BB and TL were involved in Fst enrichment analysis. IL performed bioinformatics analysis with GLP. GLP and DP were involved the RT-qPCR experiments. cDNA library construction and sequencing were performed by KL and JMA. All the authors read and approved the manuscript.

## Supporting Information

**Supplemental Information 1**: Validation of RNA-seq results by qPCR.

**Supplemental Information 2**: “Species”-responsive genes in the waterlogging and drought stress experiments.

**Supplementary Tables 1**: Overview of the cDNA libraries constructed in this study.

**Supplementary Table 2**: List of the primer pairs used for qPCR analysis.

**Supplementary Figures 1**: Schematic representation of the experimental design used in this study.

**Supplementary Figure 2** Ecophysiological characterization of the experimental design. Evolution of the O2 concentration, water content in the substrate and water potential during waterlogging or drougt stress.

**Supplementary Files 1, 2, 3, 4, 5, 6, 7, 8** are available online Inrae dataverse: G. Le Provost, B. Brachi, I. Lesur,, C. Lalanne,, K. Labadie, JM Aury, C Da Silva, D Postolache, T Leroy, C Plomion, Water stress-associated isolation barriers between two sympatric oak species , https://doi.org/10.15454/SONFVF, Portail Data INRAE, V3.0. These files includes normalized values for RNAseq data, Fold change Ratio and Gene set enrichment analysis. Supplementary file 8 include Fst value for the differentially expressed genes and their associated effect.

