## Supplementary material for "Water stress-associated isolation barriers between two sympatric oak species": Validation of RNA-seq results by qPCR.

For qPCR validation, we analyzed three genes for each experiment (waterlogging *vs.* drought) for each type of effect identified (treatment, species and interaction). Four genes displayed several different effects (Qrob_P0356680.2, Qrob_P0033250.2, Qrob_P0597160.2 and Qrob_P0475670.2), resulting in a final set of 14 unique genes. PCR amplification was unsuccessfully for five of these 14 genes, which were removed from the analysis (Qrob_P0088470.2, Qrob_P0357720.2, Qrob_P0094640.2, Qrob_P0731720.2 and Qrob_P0010280.2). A description of the genes analyzed is provided in Supplementary Table 2. For the other 10 genes, PCR efficiency ranged from 93% to 110% (Supplementary Table 2), consistent with *Taq* polymerase activity. The expression profiles of these genes are shown below in panels A (waterlogging treatment) and B (drought stress treatment).

Genes regulated during waterlogging: In total, six genes (Qrob_P0356680.2, Qrob_P0457090.2, Qrob_P0033250.2, Qrob_P0450730.2, Qrob_P0597160.2 and Qrob_P0475670.2) were validated by qPCR (Panel A). All the genes analyzed had expression profiles similar to those obtained with the RNA-seq-based approach. The first three of these genes displayed an interaction effect (Qrob_P0356680.2, Qrob_P0457090.2 and Qrob_P0450730.2). The remaining genes (Qrob_P0033250.2, Qrob_P0597160.2 and Qrob_P0475670.2) presented treatment and species effects. Qrob_P0597160.2 and Qrob_P0475670.2 were downregulated during waterlogging but overexpressed in PO. Qrob_P0033250.2 was downregulated and weakly expressed in PO, whereas it was downregulated in SO only after nine days of stress.

Genes regulated during drought stress: Four genes (Qrob_P0609950.2, Qrob_P0356680.2, Qrob_P0399130.2 and Qrob_P0612940.2) were validated by qPCR (Panel B). A similar expression profile was obtained with both approaches. Qrob_P0609950.2 displayed an interaction effect, with strong downregulation in PO and stable expression in SO during drought. Another gene (Qrob_P0399130.2) displayed a significant species effect. This gene was downregulated in both species during drought but was more strongly expressed in SO than in PO. Finally, two genes (Qrob_P0356680.2 and Qrob_P0612940.2) displayed a treatment effect only, both genes being downregulated under drought stress relative to control conditions in both species.

Panel A: qPCR validation of the candidate genes regulated during waterlogging. Abbreviations: PO: pedunculate oak, SO: sessile oak, cont: control, 24h: short-term response, 9d: long-term response. Standard deviations were obtained from three biological replicates.

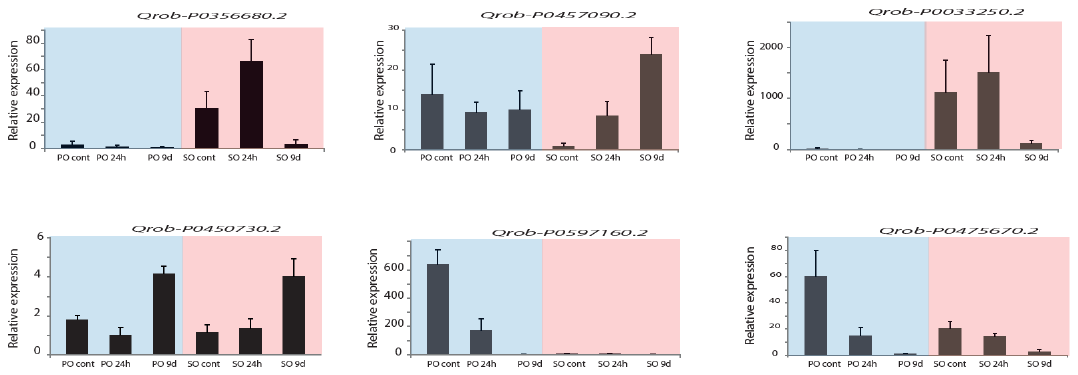

Panel B: qPCR validation of the candidate genes regulated during drought stress. Abbreviations: PO: pedunculate oak, SO: sessile oak, cont: control, ds: drought stress. Standard deviations were obtained from three biological replicates.

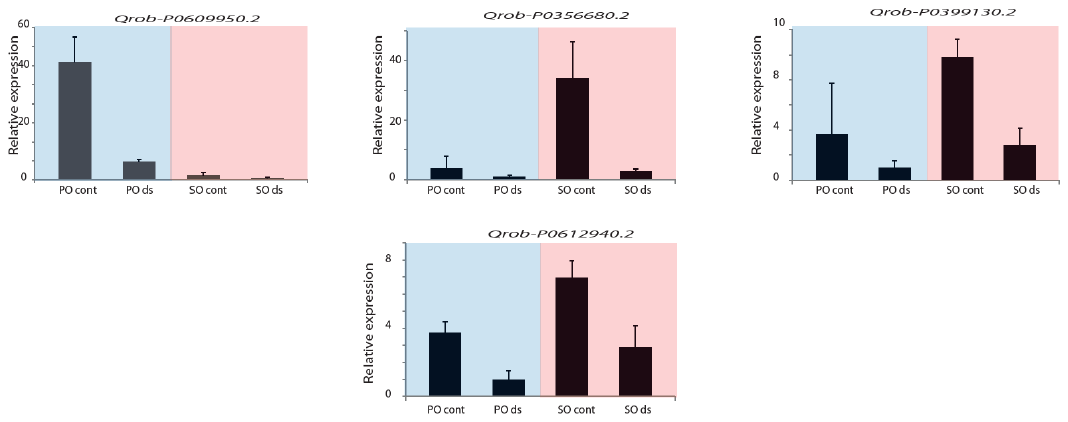
