## Supplementary material for "Water stress-associated isolation barriers between two sympatric oak species": Species responsive genes in the waterlogging and drought stress experiments.

**Results**

1/ **Gene set enrichment analysis**: All the results for gene set enrichment analysis are available in supplementary files 2 and 5, in which we show results only for the first 100 gene ontologies. A graphical representation of the first 20 ontologies is also available for a clearer view of the mechanisms regulates. The biological process (BP) GO terms with the highest degree of enrichment in PO during the waterlogging treatment were related to “brassinosteroid metabolic process”, “oxidation-reduction process”, “regulation of transcription from RNA polymerase I”, “autophagosome organization” and “suberin biosynthetic process”, whereas the metabolic function (MF) GO terms with the highest degree of enrichment concerned “nutrient reservoir activity”, “manganese ion binding”, “transcription factor activity”, “oxidoreductase activity” and “oxygen binding”. For Drought stress, we observed in SO enrichment for the following BPs:“oxidation-reduction process”, “regulation of pollen tube growth”, “brassinosteroid homeostasis”, “dimethylallyl diphosphate biosynthetic process” and “flavonoid biosynthetic process”, and for the following MFs: “iron ion binding”, “oxidoreductase activity”, “tetrapyrrole binding”, “oxygen binding” and “monooxygenase activity”).

**2/ Subnetwork enrichment analysis:** The functional networks generated with the genes displaying a species effect in the waterlogging or drought stress experiments are shown in Supplementary Figures 1 and 2 at the end of the document. For the waterlogging treatment, we identified important hubs related to vernalization, meristem identity, infection (*i.e*. plant defense and pathogen interaction), stress (drought recovery), and seed development. By contrast, for drought stress, we identified important hubs related to salinity response, ABA response and defense response. These important hubs and their associated genes are discussed below.

**Discussion**

***Waterlogging “species”-responsive genes*:** We used the 1,167 genes displaying a significant species effect for the gene-set enrichment analysis (Supplementary Figure 1 at the end of the document). We focus here on four BPs (vernalization, meristem identity, defense and seed development) comprising a large number of genes upregulated in PO (highlighted in blue in Supplementary Figure 1).

(1) Vernalization and meristem identity: We found that *AGL19* (*Qrob_P0502400.2* F_ST_<0.14) and SVP (Qrob_P0096730.2, F_ST_: 0.45) were overexpressed in PO. Both these genes encode MAD box transcription factors. Many MAD box transcription factors are specifically expressed in the root system and these factors may play a role in root development during abiotic stress responses (reviewed by Khong et al., 2008). We also identified KNAT6 (Qrob_P0640360.2, FST: 0.94), as downregulated in PO. The KNAT6 gene encodes a *KNOTTED*-like homeobox protein, which, according to Dean et al., (2004), modulates lateral root formation, its downregulation being associated with an increase in the total number of lateral roots d.

The identification of these two genes related to root growth regulation in PO suggests that anatomical modifications are of particular importance for adaptation to waterlogging in this species.

(2) Seed development: Four genes involved seed development were found to be upregulated in PO. The ABA2 gene (Qrob_P0213950.2, Fst: 0.55) encodes a xanthoxin dehydrogenase (Xu et al., 2017) involved in ABA catabolism. In cucumber, decreases in ABA levels during waterlogging are associated with an increase in the density of lateral roots, again suggesting that lateral root development is essential in hypoxic conditions. We also identified an AREB3 gene (Qrob_P0187720.2, Fst: 0.75) encoding a bZip transcription factor probably involved in ABA sensing and signaling during waterlogging, as reported by Dat et al. (2004).

The other two genes are involved in proanthocyanidin synthesis (TT2, Qrob_P0324610.2, Fst: 0.47) and oxidation (TT10, Qrob_P0691640.2, Fst: 0.52). The exact role of these genes in waterlogging adaptation is unclear, but several authors have reported that flavonoid oxidation in plants confers abiotic stress tolerance due to the scavenging of reactive oxygen species (ROS) (Pourcel et al., 2007).

(3) Defense response and pathogen interaction: A subset of the genes identified in these two hubs are related to PCD. This is not surprising, because PCD is a key component of the morphological adaptations (e.g. aerenchyma formation) observed in hypoxia-tolerant species during submergence. For example, we identified *BEN1* (Qrob_P0497760.2, Fst:0.27), *FMO1* (Qrob_P0299710.2, Fst:0.84), *BAK1* (Qrob_P0747630.2, Fst:0.35) and *RPP13* (Qrob_ P0011280.2, Fst:0.23), all of which are known to be involved in cell death. All these genes were overexpressed in PO. The *BAK1* and the *RPP13* genes encode LRR serine threonine kinases involved in the metabolism of nitric acid, a key mediator of the hypersensitive response, which involves PCD (reviewed by Breusegem et al., 2008). *BEN1* encodes an S1-type DNase responsible for the digestion of nuclear DNA during PCD (Aoyagi et al., 1998), and *FMO1* encodes a flavin-dependent monooxygenase that upregulates the ESD1 pathway known to be involved in the defense response and programmed cell death (Bartsch et al., 2006).

Finally, we identified *WRKY40* (Qrob_P0089800.2, Fst: 0.50), encoding a transcription factor overexpressed in PO. In tolerant plants, a barrier against radial oxygen loss (ROL), enhancing long-distance oxygen transport via the aerenchyma to the root tip, is formed during hypoxia. This ROL barrier is characterized by a high suberin content, suggesting that the suberin biosynthesis pathway is strongly activated during establishment of the ROL barrier. In PO, Le Provost et al. (2016) also reported that the suberin pathway is mostly expressed in hypertrophied lenticels. They hypothesized that suberin biosynthesis genes are involved in the sealing of both hypertrophied lenticels and root tips, to favor oxygen diffusion to the root system during hypoxia. Shiono et al., (2014) detected the WRKY transcription factor in the outer part of roots during the formation of a barrier to radial oxygen loss in a study based on a laser capture microdissection approach. They reported the upregulation of this transcription factor during establishment of the ROL barrier, associated with an upregulation of suberin biosynthesis genes.

Together, these results suggest that the better tolerance to waterlogging in PO results from its more efficient production of adaptive structures enhancing oxygen diffusion to the root system.

***Drought stress “species”-responsive genes***: The corresponding functional network is shown in Supplementary Figure 2 at the end of this document. Three important hubs (salinity response, ABA response and defense response) were considered particularly important. Three salinity response genes (*GER3*, *CBL10* and *PH4.6*) were upregulated in SO. *GER3* encodes a germin protein known to be downregulated during abiotic stress. Turyagyenda et al. (2013) reported that the downregulation of GER3 was associated with slower growth under water stress. The higher basal expression of *GER3* (*Qrob_P0758050.2,* F_ST_:0,28) in SO may be associated with a better capacity to maintain biomass production in drought conditions. *CBL10* (*Qrob_P0326560.2,* F_ST_:0,25) encodes a calcineurin B‐like protein involved in calcium sensing. The upregulation of this gene in *Arabidopsis thaliana* confers salt tolerance by regulating ion homeostasis (i.e. osmoregulation), conferring better tolerance to abiotic stresses, such as water deprivation (Beom‐Gi et al., 2007). PHT4.6 encodes a phosphate transporter (*Qrob_P0048620.2,* F_ST_:0.15) essential for the maintenance of primary root growth during salt stress. Cubero et al. (2009) reported that the root system was strongly reduced during water deprivation in an *Arabidopsis* mutant for this gene. Overall, the higher basal level of expression for genes involved in the maintenance of osmoregulation and primary root growth would be expected to be a major asset favoring greater drought tolerance in SO.

We also identified seven genes overexpressed in SO and involved in the ABA (abscisic acid) response (*FRO2*, *PDR12*, *K10A8.20*, *GSTF8*, *WAK2*, *CRK29* and *F9L11.8*). ABA is a key phytohormone involved in tolerance to various abiotic stresses. Its role during drought stress has been widely documented, with possible implications for the regulation of root growth during water deficit (Rowe et al., 2016). One of these genes, *WAK2* (*Qrob_P0205390.2,* F_ST_:0.50), encodes a wall-associated kinase from the RLK family. *WAK* genes may be involved in both normal cell elongation during osmotic stress and cell wall protection during dehydration (Feng et al., 2016), allowing plants to grow out of stress. *GSTF8* encodes a gluthatione-S-transferase (*Qrob_P0151730.2,* F_ST_:0.26) involved in redox homeostasis. Wang et al. (2016) reported that *GSTF8* overexpression results in an increase in glutathione conjugation, maintaining cell redox homeostasis and protecting organisms against oxidative stress. These findings suggest that GSTF8 plays a crucial role in ROS scavenging. We also identified two genes involved in ABA sensitivity (*CRK29, Qrob_P0118850.2,* Fst:0.15) or ABA transport (*PDR12, Qrob_P0561160.2,* Fst: 0.38). *CRK* genes encode cysteine-rich receptor kinases belonging to the RLK family.  Zhang et al. (2013) studied the role of CRK45 in the response to abiotic stress in *Arabidopsis thaliana*. They suggested that *CRK45* regulates the abiotic stress response by upregulating ABA responses. The *PDR12* gene (Qrob_P0561160.2 Fst: 0.23), encoding an ABC transporter involved in ABA transport, was also identified in our experiment. Kang et al., (2010) reported that a mutation of this gene in A*rabidopsis thaliana* increased drought susceptibility by affecting the uptake of ABA by cells.

Finally, we identified 11 genes (*EPR*, *PTR3*, *CYP94B3*, *RPS2*, *RPP13*, *CPR30*, *F19I3*, *MUD21.17*, *PDR12*, *RLK* and *RPM1*) related to the defense response. All but one of these genes (the exception being *RPM1*) had higher basal levels of expression in SO. Plant pathogenesis-related proteins are classically thought to be involved in plant defense and are usually induced during biotic and abiotic stresses (Haider et al., 2017), suggesting that this biological process may also underlie the better tolerance to water stress observed in SO. *CYP94B3* (*Qrob_P0369780.2,* Fst:0.22) encodes a cytochrome P450 protein. Rabara et al. (2015) reported is the involvement of this protein in jasmonoyl-isoleucine catabolism and suggested that its upregulation in tobacco may confer drought tolerance. PTR3 (*Qrob_P0351040.2,* Fst: 0.84) encodes a protein displaying similarity to a stress-induced peptide transporter. Karim et al. (2005) suggested that this protein may be involved in abiotic stress tolerance in *Arabidopsis thaliana*, through the maintenance of nitrogenous compound allocation. An *RLK* gene (*Qrob_P0495250.2,* Fst: 0.15) encoding a receptor-like kinase was also found to be overexpressed in SO. RLKs are considered to be key regulators of plant architecture and growth behavior, but they are also involved in defense and stress responses. Marshall et al. (2012) suggested that *RLK* genes may be essential for a rapid drought response and for enhanced drought tolerance.

Together, these results suggest that the better tolerance to water shortage observed in SO may be due to the overexpression of genes involved in the salinity, ABA and defense responses. These genes are involved in a key mechanism underlying drought stress tolerance and their upregulation may account for the better tolerance of water shortage observed in SO.

Supplementary Figure 1: Functional network generated with the 1,167 genes displaying a “species” effect in the waterlogging experiment. Genes highlighted in blue are upregulated in PO, whereas those highlighted in red are overexpressed in SO.

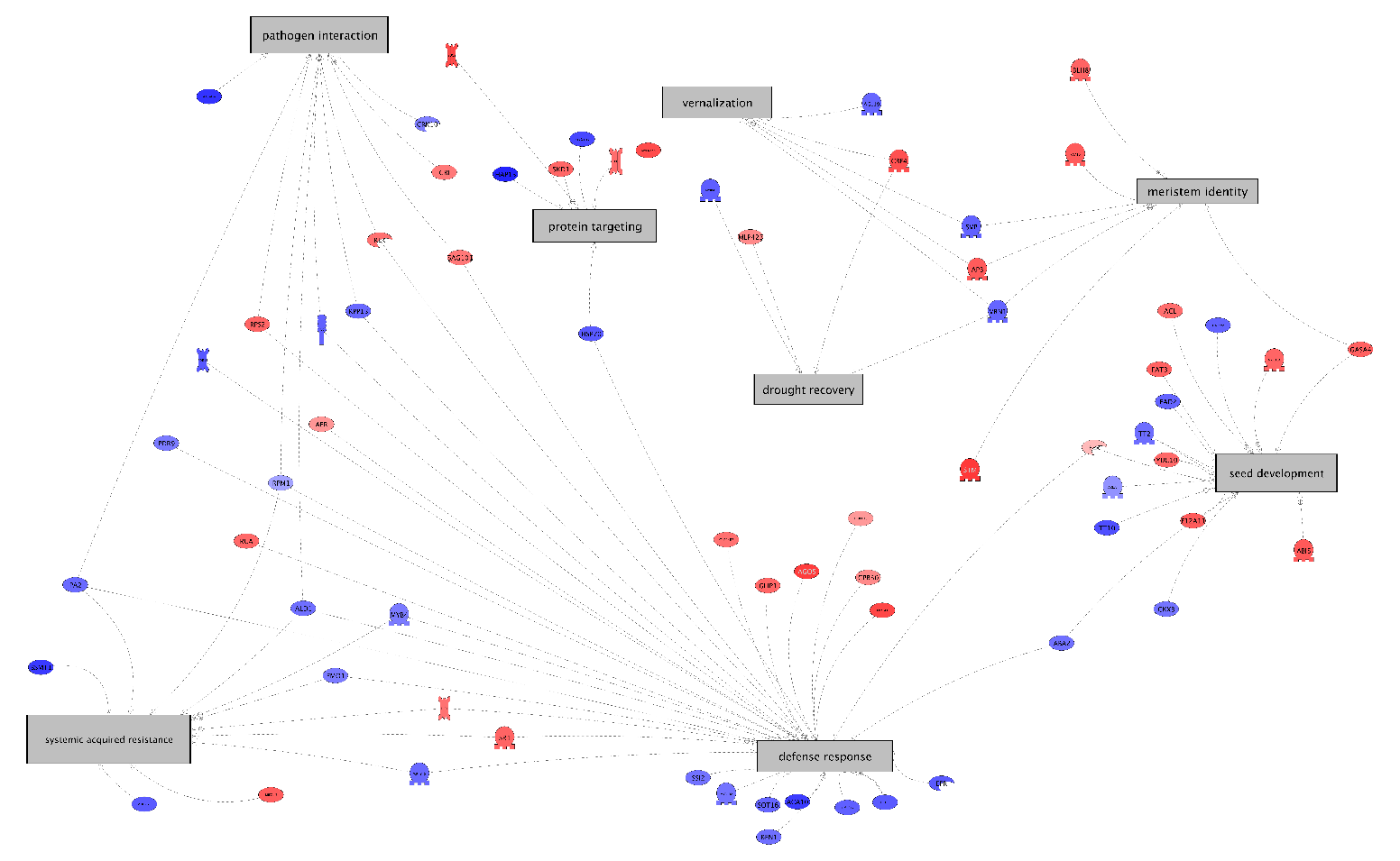

Supplementary Figure 2: Functional network generated with the 395 genes displaying a “species” effect in the drought stress experiment. Genes highlighted in red are upregulated in SO, whereas those highlighted in blue are overexpressed in PO.

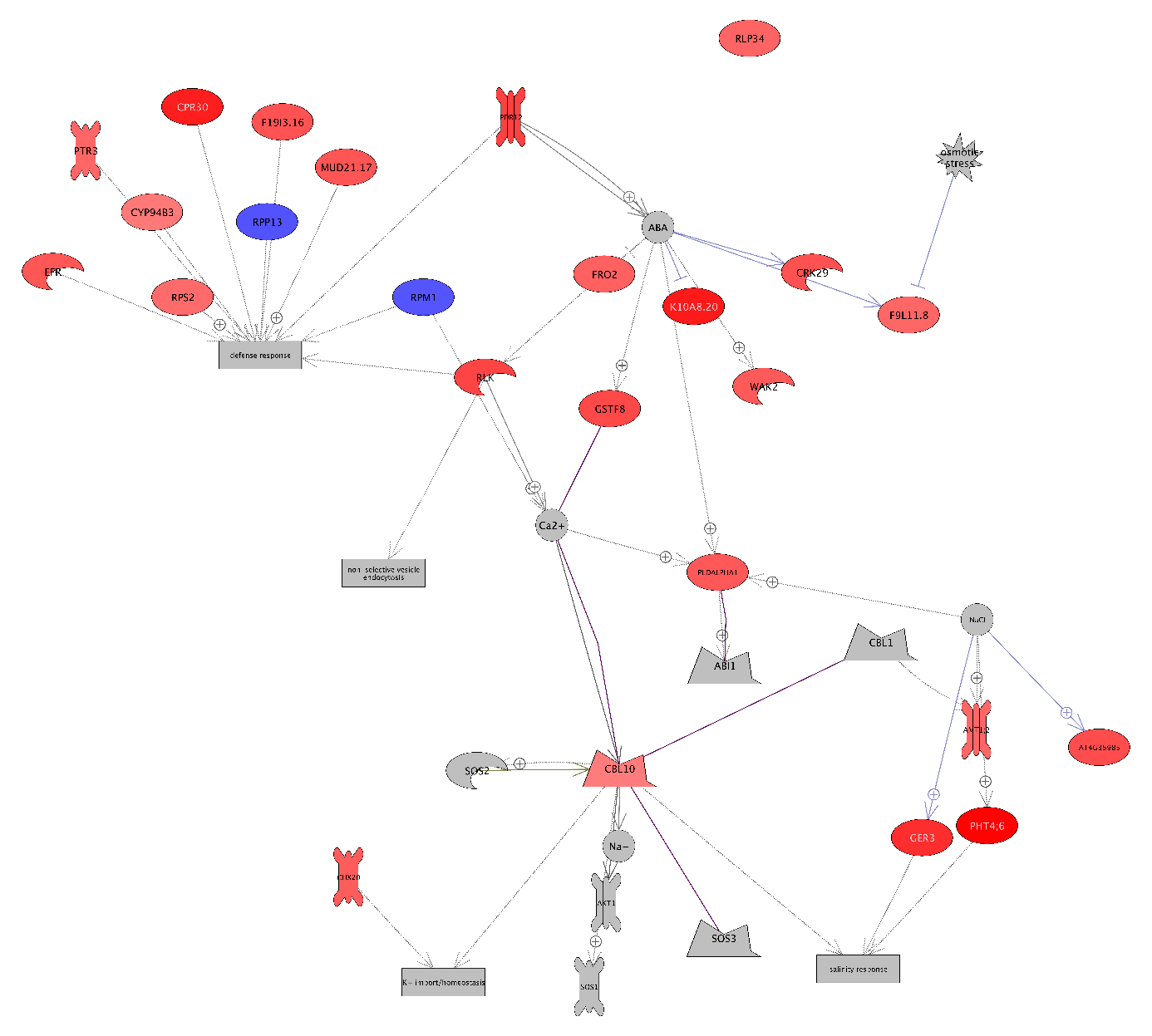
