## Supplementary material for "Water stress-associated isolation barriers between two sympatric oak species": Ecophysiological characterization of the experimental design. Evolution of the O2 concentration, water content in the substrate and water potential du

Supplementary Figure 2: Intensity of the stress applied. (A) Evolution of the O_2_ concentration in the rhizosphere along the waterlogging experiment. Standard deviation was obtained using the measurement performed on the five containers. (B) Evolution of the water content in the substrate along the drought stress treatment. (C) Evolution of the leaf predawn and the midday water potential in PO and SO respectively. Standard deviations were obtained from the measurement on five different seedlings per mother tree in both species.

**(A)**

[O2] in mg L-1

| Time | control | 5 min | 45 min | 1 hours | 2 hours | 4 hours | 6 hours | 7 hours | 1 day | 2 days | 3 days | 6 days | 7 days | 8 days | 9 days |
| --- | --- | --- | --- | --- | --- | --- | --- | --- | --- | --- | --- | --- | --- | --- | --- |
| 1 | 7.94 | 1.5 | 1.096 | 1.096 | 1.094 | 1.084 | 1.09 | 1.224 | 0.988 | 0.998 | 1.076 | 1.034 | 1.08 | 1.036 | 1.004 |

**(B)**

Mean weight of the pots (g)

| date | SO mean | PO mean |
| --- | --- | --- |
| 19 May | 519.167 | 522.833 |
| 20 May | 511.167 | 513.167 |
| 21 May | 504.25 | 504.083 |
| 22 May | 493.75 | 491.333 |
| 23 May | 451.833 | 444.5 |
| 26 May | 447.5 | 440.417 |
| 27 May | 423.417 | 416.917 |
| 30 May | 405.833 | 405.5 |
| 02 June | 405.833 | 410.909 |
| 03 June | 406.917 | 414.273 |
| 04 June | 402.833 | 410.091 |
| 05 June | 400.917 | 407.455 |
| 06 June | 368.818 | 373.455 |
| 07 June | 366.727 | 373.727 |
| 08 June | 367 | 374.364 |
| 09 June | 366.818 | 373.455 |
| 10 June | 366.636 | 373.727 |
| 11 June | 367.182 | 374.364 |
| 12 June | 366.727 | 373.455 |
| 13 June | 366.636 | 373.727 |
| 14 June | 367.182 | 374.364 |

**(C)**

**(C)**

Sessile Oaks

Water Potential (MPa)

|  | moyenne |
| --- | --- |
| Field capacity leaf predawn | 0.466667 |
| Field capacity midday | 1.642 |
| 15% of water leaf predawn | 0.923333 |
| 15% of water midday | 1.725 |

Pedunculate Oaks

|  | Water Potential (MPa) |
| --- | --- |
| Field capacity leaf predawn | 0.423333 |
| Field capacity midday | 1.205 |
| 15% of water leaf predawn | 0.9 |
| 15% of water midday | 1.62333 |

*

*

*

*
