## Supplementary material for "Water stress-associated isolation barriers between two sympatric oak species": Overview of the cDNA libraries constructed in this study.

Supplementary Table 1: Overview of the cDNA libraries constructed in this study. Abbreviations correspond to: PO: Pedunculate oak, SO: Sessile oak. The percentage of read mapped on the oak genome is indicated in parenthesis.

| **Library** | **Species** | **Biological replicate** | **Mean score**  **Phred** | **Number of reads after cleaning** | **Number of reads per condition** | **Number of reads mapped** | **SRA-NCBI Study Accession** |
| --- | --- | --- | --- | --- | --- | --- | --- |
|  | **Waterlogging Experiment** | | | | | | |
| POcont1 | PO | Rep1 (i.e mother1) | 35 | 31,082,736 | **88,854,869** | **77,051,872**  **(86%)** | PRJEB17875 |
| POcont2 | PO | Rep2 (i.e mother2) | 35 | 27,615,049 |  |  |  |
| POcont3 | PO | Rep3 (i.e mother3) | 35 | 30,157,084 |  |  |  |
| POshortterm1 | PO | Rep1 (i.e mother1) | 35 | 27,647,911 | **79,041,441** | **67,884,082**  **(85%)** |  |
| POshortterm2 | PO | Rep2 (i.e mother2) | 35 | 24,530,235 |  |  |  |
| POshortterm3 | PO | Rep3 (i.e mother3) | 35 | 26,863,295 |  |  |  |
| POlongterm1 | PO | Rep1 (i.e mother1) | 35 | 31,973,282 | **90,237,870** | **74,711,198**  **(82%)** |  |
| POlongterm2 | PO | Rep2 (i.e mother2) | 35 | 28,703,802 |  |  |  |
| Polongterm3 | PO | Rep3 (i.e mother3) | 35 | 29,560,786 |  |  |  |
| SOcont1 | SO | Rep1 (i.e mother1) | 35 | 29,135,022 | **105,448,746** | **92,037,678**  **(87%)** |  |
| SOcont2 | SO | Rep2 (i.e mother2) | 35 | 32,355,534 |  |  |  |
| SOcont3 | SO | Rep3 (i.e mother3) | 35 | 43,958,190 |  |  |  |
| SOshortterm1 | SO | Rep1 (i.e mother1) | 35 | 29,551,444 | **87,625,425** | **77,434,358**  **(88%)** |  |
| SOshortterm2 | SO | Rep2 (i.e mother2) | 35 | 31,139,081 |  |  |  |
| SOshortterm3 | SO | Rep3 (i.e mother3) | 35 | 26,934,900 |  |  |  |
| SOlongterm1 | SO | Rep1 (i.e mother1) | 35 | 27,247,656 | **85,212,082** | **71,298,544**  **(83%)** |  |
| SOlongterm2 | SO | Rep2 (i.e mother2) | 35 | 30,872,250 |  |  |  |
| Solongterm3 | SO | Rep3 (i.e mother3) | 35 | 27,092,176 |  |  |  |
|  | **Drought Stress Experiment** | | | | | | |
| POcont1 | PO | Rep1 (i.e mother1) | 35 | 27,305,022 | **86,534,414** | **73,625,692**  **(85%)** | PRJEB19536 |
| POcont2 | PO | Rep2 (i.e mother2) | 35 | 26,933,775 |  |  |  |
| POcont3 | PO | Rep3 (i.e mother3) | 35 | 32,295,617 |  |  |  |
| PO9days1 | PO | Rep1 (i.e mother1) | 35 | 24,924,521 | **74,700,596** | **62,449,658**  **(83%)** |  |
| PO9days2 | PO | Rep2 (i.e mother2) | 35 | 26,258,315 |  |  |  |
| PO9days3 | PO | Rep3 (i.e mother3) | 35 | 23,517,760 |  |  |  |
| SOcont1 | SO | Rep1 (i.e mother1) | 35 | 26,763,396 | **90,855,805** | **75,465,210**  **(83%)** |  |
| SOcont2 | SO | Rep2 (i.e mother2) | 35 | 31,804,565 |  |  |  |
| Socont3 | SO | Rep3 (i.e mother3) | 35 | 32,287,844 |  |  |  |
| SO9days1 | SO | Rep1 (i.e mother1) | 35 | 31,446,991 | **92,612,868** | **78,261,618**  **(84%)** |  |
| SO9days2 | SO | Rep2 (i.e mother2) | 35 | 30,238,729 |  |  |  |
| SO9days3 | SO | Rep3 (i.e mother3) | 35 | 30,927,148 |  |  |  |
