## Supplementary material for "Water stress-associated isolation barriers between two sympatric oak species": List of the primer pairs used for qPCR analysis.

**Supplementary Table 2**: List of the primer pairs used for qPCR analysis. Abbreviations: Tm: annealing temperature, SO: sessile Oak, PO: pedunculate Oak, ND: not differentially expressed, NA: not available, DS: Drought stress, WL: Waterlogging, int: interaction effect, treat: treatment effect, sp: species effect. Qrob_IDs were retrieved from the oak genome available in Plomion et al. (2018).

.

| Gene ID | | Significant effect in  RNAseq | Function | Forward primer(5’3’)  Reverse primer (3’-5’) | Amplicon  size (bp) | Multiband in  agarose gel | Used in  qPCR | PCR  Efficiency | Highest expression in RNAseq | Highest expression in qPCR |
| --- | --- | --- | --- | --- | --- | --- | --- | --- | --- | --- |
| Qrob_P0088470.2 | | DSint | **2-alkenal reductase (NADP(+)-dependent)** | aatgcgaccgtggaatctca  catgacttggatgcagctct | 120 | yes | No | NA | Down-regulated in SO (DS) | NA |
| Qrob_P0357720.2 | | DSint | **Hypothetical protein** | agggttggcattaggagcac  ttccaatggctcagctctgg | 115 | yes | No | NA | Up-regulated in SO(DS) | NA |
| Qrob_P0609950.2 | | DSint | **Lysine histidine transporter 1-like isoform X2** | tctcccacctcccaaacttc  atcgacaccctgctgaactc | 125 | No | Yes |  | Down-regulated in PO(DS) | Down-regulated in PO(DS) |
| Qrob_P0094640.2 | | DStreat | **Small heat shock protein** | ttggtgtagcggtcatccac  ggctcatccttacgttgctc | 114 | Yes | NO | NA | Up-regulated in PO(DS) | NA |
| Qrob_P0356680.2 | | WLint, DS treat | **Feruloyl CoA ortho-hydroxylase 3-like** | agaaggccacaatgcagagg  actcaccttccaacagcgtg | 128 | No | Yes | 99 | Down-regulated in SO (DS) | Down-regulated in SO (DS) |
| Qrob_P0399130.2 | | DSsp | **Uncharacterized protein** | gctaccaagacatcctggcc  aacccgtttcttccagaggc | 118 | No | Yes | 100 | Up-regulated in SO (DS) | Up-regulated in SO (DS) |
| Qrob_P0731720.2 | | DSsp | **Dirigent protein 21** | gctccggaatatgcgcaaag  acgaccactactttctggcc | 112 | Yes | No | NA | Down-regulated in SO | NA |
| Qrob_P0010280.2 | | DSsp | **Auxin-responsive protein** | gaggccccgagttttgtttg  ggccactgaaaccatgcaag | 116 | Yes | No | NA | Up-regulated in PO | NA |
| Qrob_P0612940.2 | | DStreat | **Hypothetical protein** | gtgctgctgagaaacttggg  ttcttctcaggtggttgggc | 115 | No | Yes | 110 | Up-regulated in SO (DS) | Up-regulated in SO (DS) |
| Qrob_P0457090.2 | | WLint | **Beta-amyrin 28-oxidase-like** | ggctggtatgggcattggat  cctggaaaagagtatgcacgg | 123 | No | Yes | 93 | Up-regulated in SO (WL) | Up-regulated in SO (WL) |
| Qrob_P0033250.2 | | WLtreat, WLsp | **Neomenthol dehydrogenase** | tcctctcgcccattcgttag  tgctaagaggacagcagagg | 121 | No | Yes | 96 | Down-regulated in SO (WL) | Down-regulated in SO (WL) |
| Qrob_P0450730.2 | | WL int | **Hypothetical protein** | tatcaccaggcccagcaatg  acgctgctaagaaccaggtc | 144 | No | Yes | 94 | Up-regulated in PO and SO (WL) | Up-regulated in PO and SO (WL) |
| Qrob_P0597160.2 | | WLtreat, WLsp | **Beta-glucosidase 13** | tgcttggaaatggggtcagg  gaccttgagccatttcgcag | 108 | No | Yes | 98 | Down-regulated in PO (WL) | Down-regulated in PO (WL) |
| Qrob_P0475670.2 | | WLtreat, WLsp | **Favonol synthase 3-like** | atctcagcctcccagtttgc  gaggccagcaccagagttag | 104 | No | Yes | 99 | Down-regulated in PO (WL) | Down-regulated in PO (WL) |
| **Control Genes** | | | | | | | | | | |
| **NA** | | Qrob_P0517760.2 | **Unknown protein** | ttgctgccttctacactggg  tggctgcattagagggacag | 102 | No | Yes | 110 | NA | NA |
| **NA** | | Qrob_P0217660.2 | **Ubiquitin-protein ligase RHF1A** | agcctatatccctggtgcca  accacgtcgcttgcatcata | 132 | No | Yes | 106 | NA | NA |
