## Supplementary material for "Water stress-associated isolation barriers between two sympatric oak species": Schematic representation of the experimental design used in this study.

### Slide 1
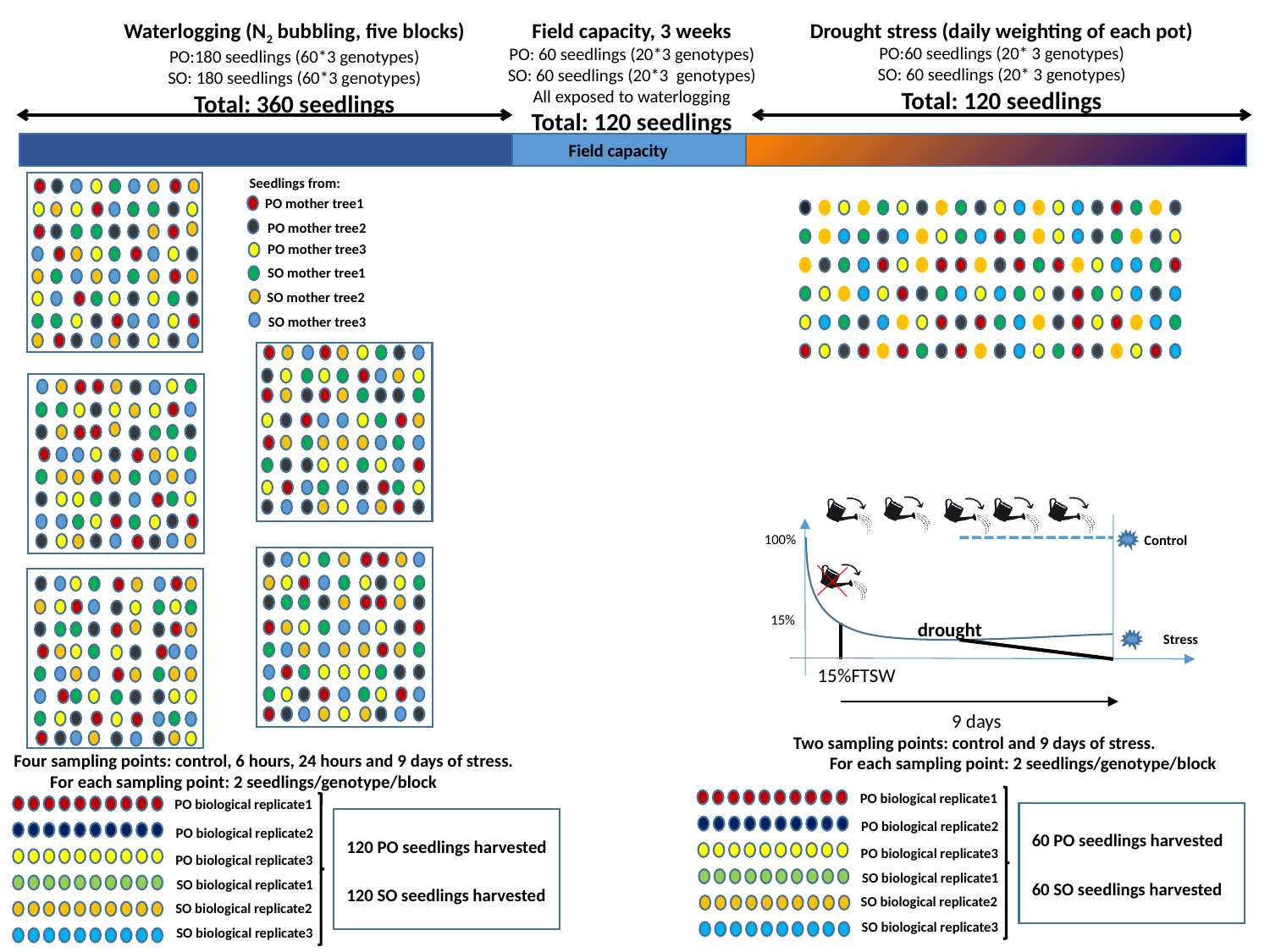

Waterlogging (N_2_ bubbling, five blocks)
PO:180 seedlings (60*3 genotypes)
SO: 180 seedlings (60*3 genotypes)
Total: 360 seedlings
Drought stress (daily weighting of each pot)
PO:60 seedlings (20* 3 genotypes)
SO: 60 seedlings (20* 3 genotypes)
Total: 120 seedlings
Field capacity, 3 weeks
PO: 60 seedlings (20*3 genotypes)
SO: 60 seedlings (20*3 genotypes)
All exposed to waterlogging
Total: 120 seedlings
Field capacity
Seedlings from:
PO mother tree1
PO mother tree2
PO mother tree3
SO mother tree1
SO mother tree2
SO mother tree3
100%
Control
15%
drought
Stress
15%FTSW
9 days
Two sampling points: control and 9 days of stress.
 For each sampling point: 2 seedlings/genotype/block
Four sampling points: control, 6 hours, 24 hours and 9 days of stress.
 For each sampling point: 2 seedlings/genotype/block
PO biological replicate1
PO biological replicate1
PO biological replicate2
PO biological replicate2
60 PO seedlings harvested
120 PO seedlings harvested
PO biological replicate3
PO biological replicate3
SO biological replicate1
SO biological replicate1
60 SO seedlings harvested
120 SO seedlings harvested
SO biological replicate2
SO biological replicate2
SO biological replicate3
SO biological replicate3
